# SETD2 negatively regulates cell size through its catalytic activity and SRI domain

**DOI:** 10.1101/2021.10.31.465891

**Authors:** Thom M. Molenaar, Eliza Mari Kwesi-Maliepaard, Joana Silva, Muddassir Malik, William J. Faller, Fred van Leeuwen

**Affiliations:** Division of Gene Regulation, Netherlands Cancer Institute, 1066CX Amsterdam, The Netherlands; Division of Oncogenomics, Netherlands Cancer Institute, 1066CX Amsterdam, The Netherlands; Department of Medical Biology, Amsterdam UMC, University of Amsterdam, 1105AZ Amsterdam, The Netherlands

**Keywords:** SETD2, histone methyltransferase, translation, cell size

## Abstract

Cell size varies between cell types but is tightly regulated by cell-intrinsic and extrinsic mechanisms. Cell-size control is important for cell function and changes in cell size are frequently observed in cancer cells. Here we uncover a non-canonical role of SETD2 in regulating cell size. SETD2 is a lysine methyltransferase and a tumor suppressor protein involved in transcription regulation, RNA processing and DNA repair. At the molecular level, SETD2 is best known for associating with RNA polymerase II through its Set2-Rbp1 interacting (SRI) domain and methylating histone H3 on lysine 36 (H3K36) during transcription. Although most of SETD2’s cellular functions have been linked to this activity, several non-histone substrates of SETD2 have recently been identified – some of which have been linked to novel functions of SETD2 beyond chromatin regulation. Using multiple, independent perturbation strategies we identify SETD2 as a negative regulator of global protein synthesis rates and cell size. We provide evidence that this function is dependent on the catalytic activity of SETD2 but independent of H3K36 methylation. Paradoxically, ectopic overexpression of a decoy SRI domain also increased cell size, suggesting that the relevant substrate is engaged by SETD2 via its SRI domain. These data add a central role of SETD2 in regulating cellular physiology and warrant further studies on separating the different functions of SETD2 in cancer development.

## Introduction

SETD2 is a lysine methyltransferase that is best known for its activity toward lysine 36 on histone H3 (H3K36), which is a histone post-translational modification found on active gene bodies (Li et al. 2016; McDaniel and Strahl 2017). H3K36 methylation by SETD2/Set2 is conserved from yeast to humans and is involved in mRNA co-transcriptional processing, repression of cryptic transcription, and DNA damage repair (Yoh et al. 2008; Luco et al. 2010; Carvalho et al. 2014; Mar et al. 2017; Huang et al. 2018). In addition, it has recently become clear that SETD2 also methylates non-histone substrates indicating that SETD2 has functions beyond chromatin regulation (Park et al. 2016; Chen et al. 2017; Seervai et al. 2020; Yuan et al. 2020). SETD2 is frequently mutated in cancer; 4.33% of all cancers carry *SETD2* mutations, with endometrial cancer, clear cell renal cell cancer, bladder cancer and colorectal cancer being most frequently associated with *SETD2* mutations (reviewed by Fahey and Davis 2017; Lu et al. 2021). Fundamental insights into the functions of SETD2 are required to understand its tumor-suppressor function.

SETD2 is capable of mono-, di- and trimethylating H3K36 *in vitro* through its catalytic SET domain. However, in cells SETD2 is only required for maintaining bulk levels of H3K36me3 but not H3K36me1/2 due to the presence of additional H3K36 mono- and dimethyltransferases in mammals (Edmunds et al. 2008; Yuan et al. 2009; Wagner and Carpenter 2012; Hyun et al. 2017; Li et al. 2019; Zaghi et al. 2020). In contrast, budding yeast only has one H3K36 methyltransferase, Set2, which is responsible for all H3K36 methylation states (Strahl et al. 2002; McDaniel and Strahl 2017). In addition to its catalytic SET domain, SETD2 contains a conserved Set2-Rbp1 interaction (SRI) domain that binds to the C-terminal domain (CTD) repeats of the largest subunit of RNA polymerase II (RNAPII) when the CTD repeats are phosphorylated at serine-2 and -5 (Sun et al. 2005). This Set2/SETD2-RNAPII interaction is essential for establishing H3K36 methylation on transcribed regions (Kizer et al. 2005; Rebehmed et al. 2014). Based on studies on Set2 in budding yeast, the emerging model is that the interaction between RNAPII and the SRI domain stimulates the activity of the catalytic SET domain rather than that it controls the localization of Set2 to active gene bodies (Youdell et al. 2008; Wang et al. 2015; Gopalakrishnan et al. 2019). Interestingly, a pathogenic point mutation observed in cancer (R2510H) in the SRI domain of human SETD2 impairs SETD2’s ability to methylate alpha-tubulin at lysine 40 during mitosis, while global methylation of H3K36 is unaffected (Park et al. 2016). Furthermore, it was recently shown that the SRI domain directly interacts with the acidic C-terminal tail of alpha-tubulin (Kearns et al. 2020). This indicates that the SRI domain not only controls the activity of SETD2 toward H3K36 but to non-histone substrates as well. It also indicates that the role of SETD2 in cancer may involve mechanisms other than defects in chromatin structure.

The lysine-specific demethylase KDM4A (also known as JMJD2A) counteracts SETD2’s function on chromatin by converting H3K36me3 into H3K36me2. In addition, KDM4A demethylates the heterochromatin mark H3K9me3. In line with the notion that many chromatin modifiers also act on non-histone proteins, KDM4A has been reported to have functions outside of the nucleus. Specifically, KDM4A associates with the initiating form of the translation machinery and stimulates mRNA translation through its catalytic activity (Van Rechem et al. 2015).

Methylation of H3K36 has two functions during transcription that are well-established in both budding yeast and mammalian cells. First, H3K36me stimulates co-transcriptional mRNA splicing by recruiting splicing factors that ‘read’ H3K36me2 or -me3 (Luco et al. 2010; Guo et al. 2014; Sorenson et al. 2016; Leung et al. 2019). Second, H3K36me2/3 promotes either the recruitment or activity of chromatin modifiers that repress (cryptic) transcription initiation from within actively transcribed gene bodies (Carrozza et al. 2005; Keogh et al. 2005; Lickwar et al. 2009; Joshi and Struhl 2005; Baubec et al. 2015; Neri et al. 2017). Another potential function of H3K36 methylation is to promote histone recycling during transcription elongation. Nucleosomes act as barriers for transcription and are therefore transiently disrupted to allow passage of RNAPII (Bondarenko et al. 2006; Petesch and Lis 2012; Studitsky et al. 2016; Chen et al. 2019). In the wake of transcription, histones can either be recycled or replaced by newly synthesized histones, leading to histone turnover. In budding yeast, Set2 represses histone turnover in active genes indicating that Set2 promotes histone recycling during transcription (Venkatesh et al. 2012; Smolle et al. 2012; Radman-Livaja et al. 2012). It is currently unclear if SETD2 has a similar function in mammalian cells. Interestingly, SETD2 promotes both the localization of the conserved histone chaperone FACT (<u>fa</u>cilitates chromatin transcription) to chromatin as well as the maintenance of proper nucleosome organization in active genes in human cells (Carvalho et al. 2013; Simon et al. 2014). Given that FACT promotes histone recycling during transcription in budding yeast (Jamai et al. 2009; Jeronimo et al. 2019) and in *in vitro* studies (Hsieh et al. 2013; Farnung et al. 2021), an attractive model is that SETD2-mediated recruitment of elongation factors such as FACT maintains chromatin integrity (i.e. nucleosome occupancy) during transcription.

Here, we set out to investigate SETD2’s role in maintaining histone levels. We found that depletion of SETD2 alters the ratio between cellular protein content and histone proteins. This altered histone over total protein ratio was not due to a loss of chromatin integrity leading to global loss of histones from DNA but rather due to an increase in total cellular protein content and cell size. Protein content is controlled by protein synthesis and degradation rates, and can be coordinated at the level of both transcription as well as translation. Mechanistically, we demonstrate that SETD2 controls global protein synthesis rates, and we provide evidence that this function is dependent on SETD2 catalytic activity and the SRI domain but most likely independent of H3K36me3. Our results suggest that SETD2 acts opposite to the demethylase KDM4A (Van Rechem et al. 2015) to regulate protein synthesis and cell size.

## Results

### SETD2 controls total protein content

In metazoans, compromised chromatin integrity leads to the deposition of the replication-independent histone variant H3.3. H3.3 acts as a ‘gap-filler’ histone and prevents the accumulation of naked DNA when histone deposition (e.g. during DNA replication) is compromised (Ray-Gallet et al. 2002; Tagami et al. 2004; Maze et al. 2015; Tvardovskiy et al. 2017). Our initial aim in this study was to determine if SETD2 represses the deposition of the H3.3 gap filler histone, given that (1) Set2 represses replication-independent histone turnover in active genes in budding yeast (Venkatesh et al. 2012) and (2) SETD2 maintains nucleosome occupancy in active gene bodies (Carvalho et al. 2013; Simon et al. 2014). We therefore depleted SETD2 in human retinal pigment epithelial cells transduced with the telomerase gene (RPE1-hTERT), which is a non-transformed near diploid human cell line (designated RPE1 from here on). To monitor H3.3 (which differs five amino acids from H3.1 and four amino acids from H3.2), we used RPE1 cells carrying a endogenously V5 epitope-tagged copy of the H3.3 gene *H3F3B* (Molenaar et al. 2020). Despite being frequently inactivated in cancer, SETD2 is an essential gene in several human cell lines (Blomen et al. 2015; Wang et al. 2015; Bertomeu et al. 2018). Therefore, to prevent looking at potential secondary effects of long term SETD2 loss, we employed an inducible SETD2 knockdown system using doxycycline (dox) inducible miRNAs against *SETD2* based on the miR-E optimized backbone (Fellmann et al. 2013).

Treating RPE1-*H3F3B*-V5 cells transduced with inducible miRNAs targeting *SETD2* with dox for 72h led to a reduction in *SETD2* mRNA expression (**Supplementary Figure 1A**) and H3K36me3 levels (**Figure 1A, B**), as expected. We first assessed global H3.3-V5 levels in protein-normalized whole-cell lysates from SETD2 depleted RPE1 cells. Unexpectedly, H3.3 levels were significantly reduced in SETD2 depleted cells (**Figure 1A, B**). This was unexpected for two reasons. First, we predicted that SETD2 *represses* H3.3 deposition in active gene bodies. Second, only a small percentage of the human genome constitutes active gene bodies (i.e. only 1-5% of nucleosomes is marked by H3K36me3; LeRoy et al. 2013) and we therefore did not expect global changes in H3.3 levels upon SETD2 depletion. Strikingly, in addition to H3.3, we also observed that protein-normalized whole-cell lysates from SETD2 depleted cells had reduced histone H3 and H4 levels compared to untreated cells or cells expressing a scrambled miRNA (**Figure 1A**). Does SETD2 maintain global histone levels (i.e. chromatin integrity) or does SETD2 maintain a normal DNA to total protein ratio? To answer this, we measured genomic DNA levels by qPCR in protein-normalized cell lysates and found that SETD2 depleted lysates had lower DNA levels (**Figure 1A** lower panel). This suggests that the DNA:protein ratio is lowered by SETD2 loss, and that histones appropriately scale with DNA levels in SETD2 knockdown cells. Indeed, when normalizing protein lysates for genomic DNA levels (which equals normalizing for cell numbers), SETD2 depleted cells showed similar histone levels and increased levels of non-histone proteins such as α-tubulin and β-actin (**Figure 1C**). This suggests that SETD2-depleted cells have an increased total cellular protein content.

**Figure 1.**
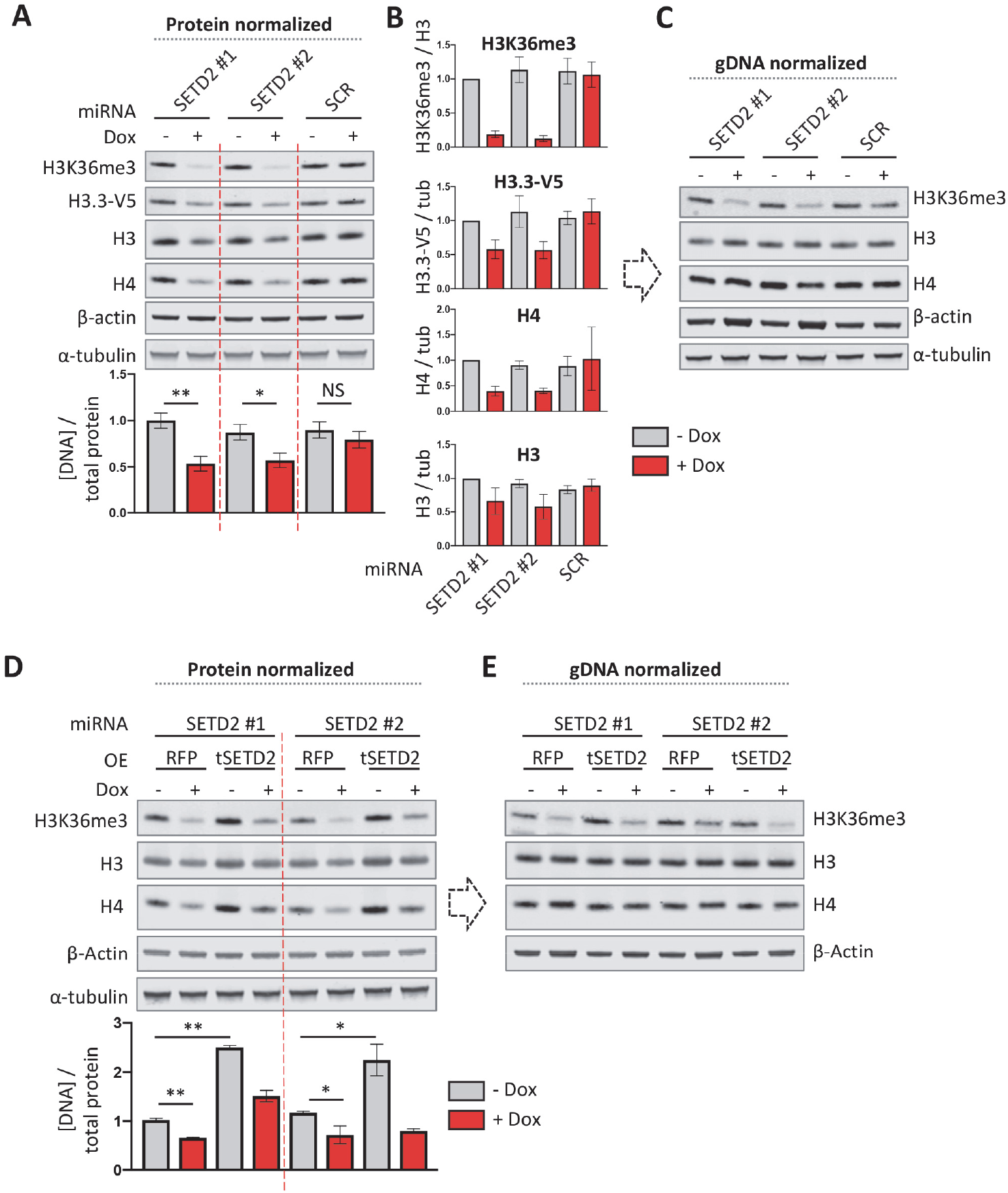
SETD2 depletion increases the total protein content of human cells. (**A**) Western blot of RPE1 cells with doxycycline-inducible knockdown of *SETD2*. Cells were treated with doxycycline for 72h. Cell lysates were normalized for total protein (left panel) or genomic DNA content (right panel). The bar plot below the left panel represents genomic DNA levels quantified by qPCR in protein normalized lysates. (**B**) Western blot of RPE1 cells with doxycycline-inducible knockdown of *SETD2* (72h induction) and constitutive overexpression of either RFP (control) or N-terminally truncated SETD2 (tSETD2). Cell lysates were normalized for total protein (left panel) or genomic DNA content (right panel). The bar plot below the left panel represents genomic DNA levels quantified by qPCR in protein normalized lysates. (**C**) 2D cell size as measured by image flow-cytometry of RPE1 cells with inducible SETD2 depletion and/or constitutive tSETD2 overexpression (RFP as control). Cells were treated with doxycycline for 72h for inducible miRNA based *SETD2* knockdown. SCR, scramble miRNA; OE, overexpression; Dox, doxycycline. Error bars represent SD of three biological replicates.

To confirm that this phenotype was indeed caused by loss of SETD2 expression, we determined if the increased protein content in SETD2-depleted cells could be rescued by overexpressing miRNA-resistant SETD2. We used a catalytically active but truncated version of SETD2 (tSETD2) that lacks the first 504 amino acids of the unstructured N-terminal domain to facilitate expression (Carvalho et al. 2013). Interestingly, tSETD2 overexpression increased the DNA / protein ratio, indicating that total cellular protein content was reduced in these cells (**Figure 1D, E**). Combining tSETD2 overexpression and endogenous SETD2 knockdown restored protein content to approximately wild-type levels.

In addition to miRNA-based knockdowns, we suppressed SETD2 expression using an independent alternative approach, CasRx (Cas13d) mediated RNA cleavage. The CasRx system has been reported to have a high knockdown efficiency with minimal off-target effects in human cells (Konermann et al. 2018). Indeed, we observed high mRNA cleavage efficiency using two *SETD2* mRNA targeting guide RNAs (gRNAs). Importantly, CasRx-mediated knockdown of SETD2 also led to an increase in total protein content, confirming the miRNA-based SETD2 knockdown results (**Supplementary Figure 1B**). Furthermore, to determine if increased protein content upon SETD2 depletion was restricted to RPE1 cells, we knocked down SETD2 in two other normal human cell lines: the human fetal lung fibroblast cell line TIG3 and the foreskin fibroblast cell line BJET. We observed a decrease in histone H3 and H4 in protein-normalized lysates from SETD2 depleted TIG3 and BJET cells (**Supplementary Figure 1C**) indicating that SETD2 controls total cellular protein levels in multiple human cell lines.

### SETD2 controls protein synthesis rates and cell size

Global protein levels and cell size are closely correlated. Therefore, the observed protein content regulation by SETD2 should presumably lead to an alteration in cell size as well. Indeed, we observed by imaging flow-cytometry that SETD2 depletion increased cell size (measured as 2D cell surface of cells in suspension) while tSETD2 overexpression decreased cell size (**Figure 2A**). Taken together, these results suggest that SETD2 controls total protein content and consequently cell size.

**Figure 2.**
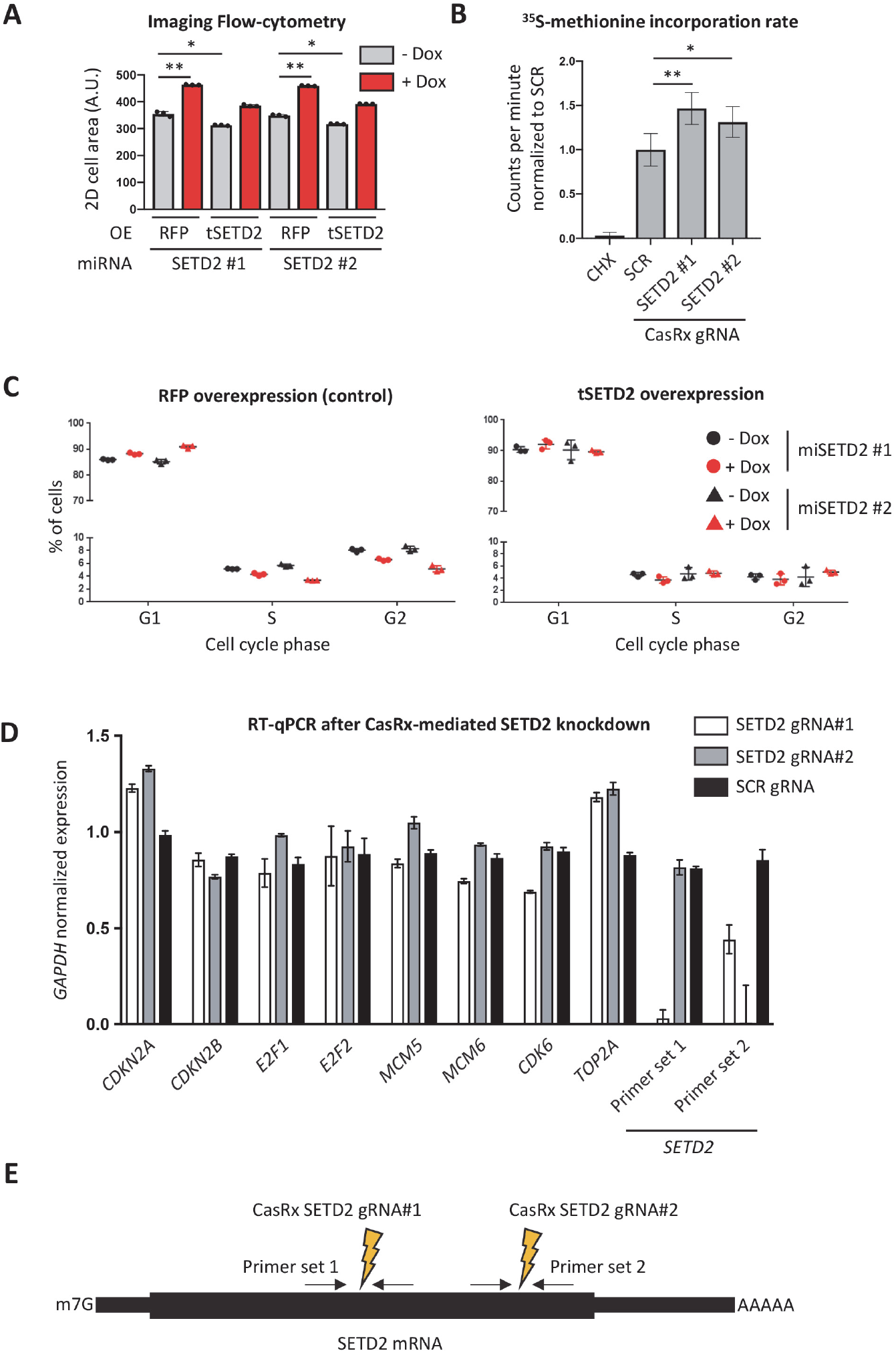
Inducible depletion of SETD2 increases protein synthesis rates accompanied with a minor accumulation of cells in G_1_. (**A**) ^35^S-methionine incorporation assay of RPE1 cells 72h following doxycycline-induced CasRx-based *SETD2* knockdown. CHX indicates a control experiment in which cells were treated with cycloheximide for 1h, which inhibits protein synthesis. (**B**) Cell cycle distribution as measured by flow-cytometry of propidium iodide stained SETD2 depleted and tSETD2 overexpressing RPE1 cells. Cells were treated with doxycycline for 72h for inducible expression of *SETD2* targeting miRNAs, while RFP (control; left panel) and tSETD2 overexpression (right panel) was constitutive. (**C**) RT-qPCR for mRNA expression analysis of genes involved in cell cycle regulation in RPE1 cells, 72h following doxycycline-induced CasRx-based *SETD2* knockdown. (**D**) The two CasRx gRNAs used for targeting *SETD2* mRNA are each flanked by a RT-qPCR primer pair used for *SETD2* expression analysis in (C). Error bars represent SD of three biological replicates.

An increased cell size can be accompanied by adaptations in protein synthesis and degradation, two opposing but coupled processes. To directly measure protein synthesis rates in SETD2 depleted cells, we used a radioactively labeled ^35^S-methionine incorporation assay. SETD2 depletion using the CasRx system led to a significant increase in the incorporation rate of ^35^S -methionine normalized for total protein content (**Figure 2B**). This indicates that SETD2 negatively regulates protein synthesis and suggests that the increased protein content in SETD2 depleted cells is caused by an increase in protein synthesis rate.

Mammalian cells that are arrested in G1 and exposed to growth factors generally continue to increase in cell size and have an increased protein synthesis rate compared to proliferating cells (Conlon and Raff 2003). We therefore used cell cycle profiling by flow-cytometry to determine if inducible SETD2 depletion led to a G1 arrest. SETD2 depletion led to a slight increase in the number of cells in G1 and a decrease in the number of cells in S phase and G2 (**Figure 2C**). tSETD2 overexpression also led to a small increase in the number of cells in G1 (compare between left and right plots). However, knockdown of endogenous SETD2 did not rescue cell cycle distribution in tSETD2 expressing cells, even though SETD2 knockdown did partially restore the size of tSETD2 expressing cells as measured by imaging flow-cytometry (see **Figure 2A**). To look at cell cycle defects in SETD2 depleted cells in an independent way, we measured the mRNA levels of several genes involved in cell cycle progression in RPE1 cells in which SETD2 was depleted using the dox-inducible CasRx system (**as in Supplementary Figure 1B**). Consistent with the cell cycle distribution analysis, SETD2 depletion resulted in a minor decrease in the expression of genes involved in cell cycle progression such as *E2F1/2, MCM5/6* and *CDK6* (**Figure 2D, E**). However, it seems unlikely that the small difference in cell cycle distribution is the primary reason for the increased cell size in SETD2 depleted cells.

### SETD2 controls cell size through its catalytic activity

How does SETD2 control cell size? We first wanted to determine if SETD2 controls cell size through its catalytic activity. However, we were unable to establish RPE1 cell lines stably (over)expressing catalytically inactive tSETD2 (tSETD2-Q1669A) suggesting that this is lethal in RPE1 cells. As an alternative approach to determine the role of SETD2’s catalytic activity in regulating cell size, we stably overexpressed the demethylase KDM4A in RPE1 cells. KDM4A (also known as JMJD2A) counteracts SETD2’s function on chromatin by converting H3K36me3 into H3K36me2. In addition, KDM4A demethylates the heterochromatin mark H3K9me3. Stable KDM4A overexpression decreased global H3K36me3 and H3K9me3 levels in RPE1 cells, as expected (**Figure 3A**). Importantly, KDM4A overexpression increased the total cellular protein content similar to SETD2 depletion (**Figure 3A**). This result suggests that SETD2 controls protein content through its catalytic activity and opens up the possibility that SETD2 and KDM4A act in the same pathway.

**Figure 3.**
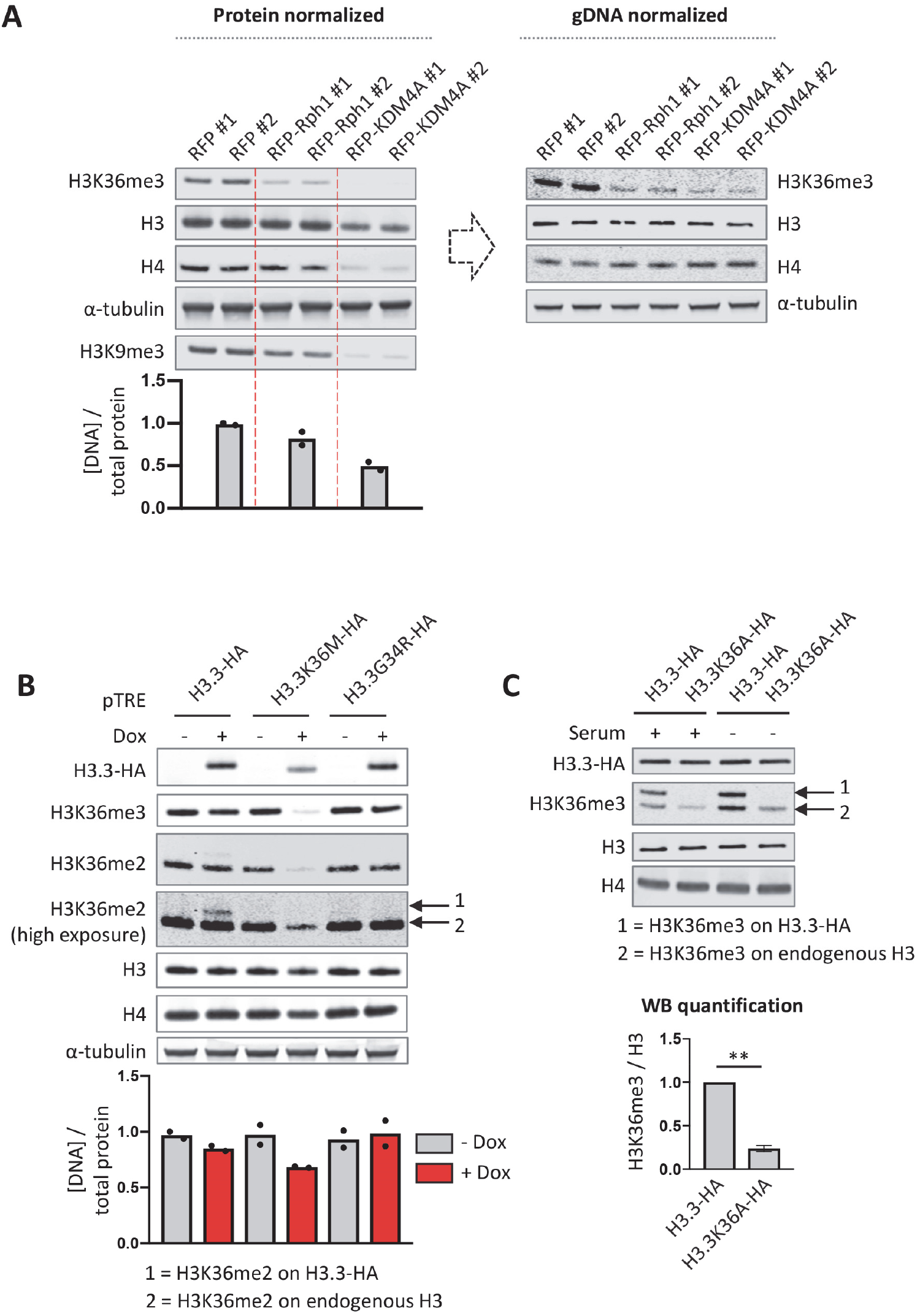
SETD2 controls cellular protein content through its catalytic activity. (**A**) Western blot of RPE1 cells constitutively overexpressing the yeast demethylase Rph1 or human H3K36me3/H3K9me3 demethylase KDM4A. Cell lysates were normalized for total protein (left panel) or genomic DNA content (right panel). The bar plot below the left panel represents genomic DNA levels quantified by qPCR in protein normalized lysates. (**B**) Western blot of RPE1 cells with doxycycline inducible overexpression of hemagglutinin (HA) epitope-tagged “onco” H3.3 histones. The bar plot represents genomic DNA levels quantified by qPCR in protein normalized lysates. (**C**) Western blot of RPE1 cells with stable overexpression of H3.3-HA and H3.3K36A-HA. Dots represent the individual values of two biological replicates (in A and B). Error bars represent SD of three biological replicates (in C).

In line with the notion that many chromatin modifiers also act on non-histone proteins, KDM4A has been reported to have functions outside of the nucleus. Specifically, KDM4A associates with the initiating form of the translation machinery and stimulates protein synthesis rates through its catalytic activity (Van Rechem et al. 2015). This suggests that KDM4A mediated demethylation of a component of the translation machinery stimulates protein synthesis. However, the identity of this methylated substrate and the methyltransferase involved are unknown. SETD2 is best known for its ability to methylate H3K36. However, the list of non-histone substrates that are methylated by SETD2 continues to grow. In an attempt to determine if SETD2/KDM4A regulate protein synthesis via H3K36, we also overexpressed the budding yeast homologue of KDM4A, Rph1 (<u>R</u>egulator of <u>PH</u>R1) which demethylates both H3K36me2 and H3K36me3 in *Saccharomyces cerevisiae* (Kim and Buratowski 2007; Klose et al. 2007). Stable Rph1 overexpression in RPE1 cells decreased H3K36me3 levels and had a small effect on H3K9me3 (which is absent in *S. cerevisiae*) but did not significantly alter total cellular protein content (**Figure 3A**). One possible explanation for the differential effects between Rph1 and KDM4A overexpression is the they both act on H3K36 but have likely evolved in conjunction with the opposing methyltransferases (here Set2 and SETD2) to act on additional species-specific substrates. Taken together, these results suggest that SETD2 regulates cell size through its methylation activity but argue against direct involvement of its activity toward H3K36.

To further corroborate these findings, we inhibited SETD2 function by overexpressing the H3.3K36M oncohistone. H3.3K36M, a mutant histone found in chondroblastoma (Behjati et al. 2013), inhibits SETD2 as well as the H3K36 mono-and dimethyltransferase NSD2, in a dominant negative manner i.e. *in cis* and *in trans* (Lewis et al. 2013; Lu et al. 2016; Zhang et al. 2017). As a control, we also overexpressed H3.3G34R which is found in glioblastoma (Schwartzentruber et al. 2012) and osteosarcoma (Behjati et al. 2013) and which inhibits SETD2 only locally *in cis* (Fang et al. 2018; Shi et al. 2018). Inducible overexpression of HA-tagged H3.3K36M but not H3.3G34R reduced global H3K36me2 and H3K36me3 levels, as expected (**Figure 3B**). Interestingly, H3.3K36M overexpression lowered the genomic DNA:protein ratio but not to the same extent as SETD2 depletion or KDM4A overexpression, despite H3K36me3 being almost completely absent. This shows that there is no direct correlation between global H3K36me3 levels and cell size. However, it cannot be excluded that the remaining H3K36me3 localized on a specific set of genes and indirectly regulates protein content.

To more directly investigate the involvement of H3K36 methylation in regulating cell size, we stably overexpressed H3.3 or H3.3K36A in RPE1 cells with the aim to replace a substantial fraction of H3.3 (and canonical H3) with an H3.3K36A histone mutant that cannot be methylated on K36. Humans have 15 genes encoding H3 and H3.3, making it difficult to assess the function of histone modifications by mutating endogenous H3 amino acid residues, a strategy that has been successfully employed in yeast and flies (Meers et al. 2017). Based on H3K36me3 immunoblotting, we found that ectopic expression by the strong *EEF1A1* promoter led to high incorporation levels of ectopic HA-tagged H3.3 (H3.3-HA). Note that the C-terminal HA tag interferes with the recognition of the anti-H3 antibody (Abcam 1791). Since H3.3 accumulates in non-dividing cells (Maze et al. 2015), we also attempted to further increase the level of ectopic H3.3-HA incorporation by depriving RPE1 cells of serum. However, we found that 7 days of serum deprivation did not lead to higher levels of H3.3-HA in RPE1 cells. H3.3K36A has a minor *trans* inhibitory effect on SETD2 although not as strong as H3.3K36M (Lu et al. 2016). Indeed, we observed that high expression of H3.3K36A-HA (which cannot be methylated and is not recognized by the H3K36me antibodies) reduced the levels H3K36me3 on endogenous histone H3 (**Figure 3C**). H3.3K36A overexpression did not affect cell size, despite total H3K36me3 levels (i.e. on both ectopic and endogenous H3) being significantly reduced. This provides further support for the model that SETD2 controls cell size independently of H3K36me3.

### SETD2 controls cell size through its SRI domain

To gain further mechanistic insight into how SETD2 negatively regulates cell size, we targeted the interaction between SETD2 and RNAPII. This interaction is mediated by the SRI domain, which is conserved from yeast Set2 to human SETD2. The SRI domain interacts with the CTD of RNAPII when phosphorylated at serine 2 and serine 5 in the heptapeptide repeat and this interaction is essential for establishing H3K36me3 in both yeast and human cells (Kizer et al. 2005; Sun et al. 2005; Rebehmed et al. 2014). Ectopic overexpression of the *S. cerevisiae* Set2 SRI domain (SRI_Set2_) fused to a nuclear localization signal (NLS) reduced global H3K36me3 levels in RPE1 cells, presumably because the excess free SRI_Set2_ domain acts as a decoy for RNAPII (**Figure 4A**). Importantly, SRI_Set2_ overexpression increased cell size (**Figure 4B**). This indicates that SETD2 regulates cell size through its SRI domain. To determine if the nuclear localization of this decoy SRI_Set2_ domain was important for its ability to disrupt cell size regulation, we also overexpressed a SRI_Set2_ domain fused to the HIV Rev protein nuclear export signal (NES). However, we were unable to generate RPE1 cell lines stably overexpressing NES-SRI_Set2_, suggesting that this is toxic in RPE1 cells.

**Figure 4.**
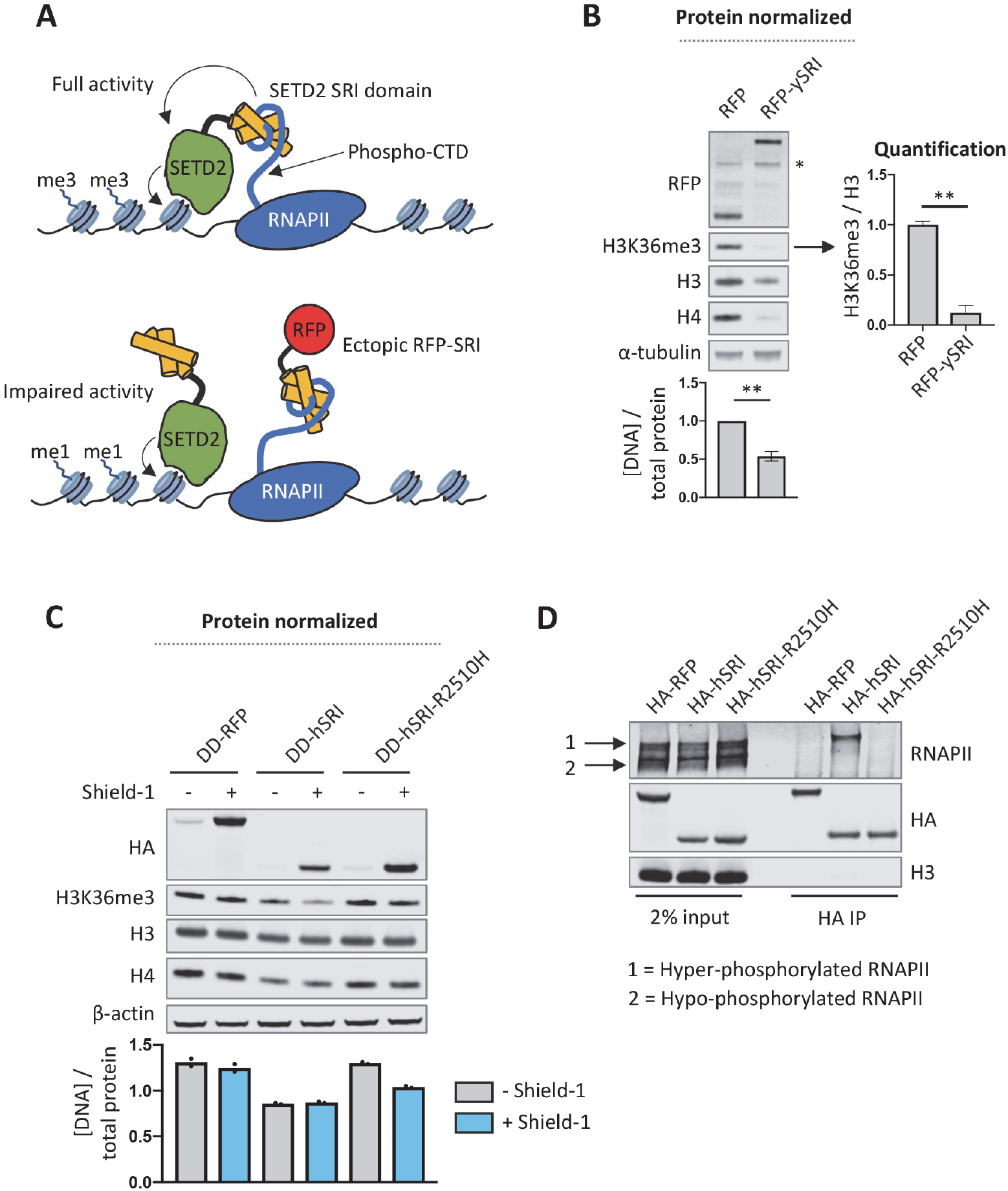
Ectopic overexpression of the Set2/SETD2 SRI domain inhibits H3K36me3 and increases cellular protein content. (**A**) Cartoon to illustrate how overexpression of a “decoy” SRI domain might specifically interrupt SETD2 activity toward H3K36 (as well as other SRI-dependent SETD2 substrates). (**B**) Western blot of RPE1 cells stably overexpressing the yeast Set2 SRI domain N-terminally fused to tagRFP and an SV40 nuclear localization signal (NLS). The bar plot represents genomic DNA levels quantified by qPCR in protein normalized lysates. (**C**) Western blot of RPE1 cells stably overexpressing the human SETD2 SRI domain N-terminally tagged with a destabilizing domain (DD; stabilized by Shield-1), HA-tag and SV40 NLS. DD-RFP is tagRFP N-terminally fused to DD-HA-SV40 NLS. The bar plot represents genomic DNA levels quantified by qPCR in protein normalized lysates. (**D**) Western blot of the ectopically overexpressed SETD2 SRI domains immunoprecipitated from HEK293T cells. HEK293T cells were transiently transfected with DD-HA-NLS-tagRFP (control), DD-HA-NLS-SRI or DD-HA-NLS-SRI-R2510H encoding plasmids and treated with Shield-1. Cells were lysed 48h after transfection, and RFP or SRI domains were immunoprecipitated with anti-HA antibody. Dots represent the individual values of two biological replicates (in C). Error bars represent SD of three biological replicates (in B).

Although the SRI domain is best known for its ability to interact with the RNAPII phospho-CTD, the SRI domain also contributes to the ability of SETD2 to methylate non-histone substrates. For example, a pathogenic point mutation in the SRI domain of SETD2 (R2510H) disrupts alpha-tubulin K40 methylation by SETD2 (Park et al. 2016). In line with this, the SETD2 SRI domain was recently shown to directly interact with the C-terminal tail of alpha-tubulin (Kearns et al. 2020). Interestingly, while mutating the R2510 residue in the SETD2 SRI domain to an alanine (R2510A) disrupts the interaction between the SRI domain and RNAPII (Li et al. 2005), the R2510H mutation disrupts alpha-tubulin K40 methylation but not H3K36 methylation (Park et al. 2016). Therefore, SRI_SETD2_-R2510H can be used to functionally separate SETD2-mediated alpha-tubulin methylation from RNAPII-mediated H3K36 methylation.

To strengthen the finding that SRI_Set2_ overexpression disrupts SETD2-mediated cell size regulation, we established RPE1 cells stably expressing human SRI_SETD2_ tagged with an NLS and a destabilizing domain (DD) that allows for Shield-1 inducible protein expression (Banaszynski et al. 2006). Similar to SRI_Set2_ overexpression, SRI_SETD2_ reduced H3K36me3 levels when expressed at high levels (i.e. stabilized by Shield-1) and increased cellular protein content (**Figure 4C**). Interestingly, at lower expression levels (i.e. without Shield-1) SRI_SETD2_ did not strongly affect global H3K36me3 levels while it still increased protein content compared to RFP expressing control cells. This provides further evidence that H3K36me3 and protein content regulation by SETD2 are decoupled from each other. We also established RPE1 cells stably expressing SRI_SETD2_-R2510H. Surprisingly, SRI_SETD2_-R2510H overexpression did not affect H3K36me3 levels and only slightly increased protein content when stabilized by Shield-1, despite being expressed at somewhat higher levels than SRI_SETD2_. Although the interaction between SETD2 and RNAPII is required for H3K36me3 and SETD2-R2510H can still establish H3K36me3 (Hacker et al. 2016; Park et al. 2016) our findings suggest that when overexpressed the R2510H mutation abolishes the function of SRI_SETD2_ as a decoy for RNAPII. To test this assumption, we immunoprecipitated the ectopic SRI domains from transiently transfected HEK293T cells and found that SRI_SETD2_ but not SRI_SETD2_-R2510H interacted with RNAPII (**Figure 4D**). This lack of interaction between SRI_SETD2_-R2510H and RNAPII explains why SRI_SETD2_-R2510H does not reduce H3K36me3 levels upon overexpression, as it likely does not outcompete endogenous SETD2 for RNAPII binding. Collectively, these results demonstrate that SETD2 regulates protein content by engaging a substrate through its SRI domain, and that the R2510 residue in SRI is essential for this interaction.

## Discussion

SETD2 has multiple cellular functions including RNA processing, the repression of cryptic transcription, and DNA repair (Yoh et al. 2008; Luco et al. 2010; Carvalho et al. 2014; Mar et al. 2017; Huang et al. 2018). Mechanistically, most of these processes have been shown to involve the classic molecular function of SETD2, i.e. H3K36 methylation. However, as additional non-histone SETD2 substrates continue to be identified it is becoming clear that SETD2’s function extends beyond chromatin and transcription regulation (Park et al. 2016; Chen et al. 2017; Seervai et al. 2020; Yuan et al. 2020). Here, we report a novel cellular function of SETD2, namely the regulation of protein synthesis rate and cell size. We showed that SETD2 exerts this function through its catalytic activity as overexpression of the demethylase KDM4A has a similar phenotype as SETD2 depletion. Our results are consistent with the previously reported findings that KDM4A stimulates protein synthesis (Van Rechem et al. 2015). However, we cannot exclude at this point that SETD2 inhibits mRNA translation through a pathway that is independent of KDM4A. Protein synthesis takes up a large proportion of the energy available to a cell and is therefore tightly regulated by a wealth of mechanisms. It remains to be determined if the increased translation rates in SETD2 depleted cells are an indirect consequence of for example deregulated signaling pathways or cell cycle control, or if SETD2 controls translation in a more direct way, perhaps in concert with KDM4A.

An important step to determine the mechanism through which SETD2 regulates cell size will be to identify the relevant substrate methylated by SETD2. H3K36me3 is the classical SETD2 substrate and could conceivably regulate the expression of genes involved in translation for example by regulating mRNA splicing (Luco et al. 2010; Simon et al. 2014; Leung et al. 2019). However, several lines of evidence suggest that SETD2 regulates translation independently of H3K36me3. First, unlike SETD2 depletion, overexpression of the yeast demethylase Rph1 did not affect total cellular protein content. However, because H3K36me3 was not completely abolished in these cells it is possible that local H3K36me3 on certain genes is sufficient to maintain normal protein content. A similar argument could be made for H3.3K36A overexpression, which did not affect protein content but also did not completely remove H3K36me3 on endogenous H3. Second, overexpression of the H3.3K36M oncohistone almost completely removed H3K36me3 but did not affect protein content as strongly as SETD2 knockdown. This suggests that there is no direct correlation between SETD2 activity toward H3K36 and protein content. H3.3K36M acts by inhibiting SETD2 activity *in cis and in trans* but it is not completely understood if H3.3K36M inhibits all SETD2 protein or only the SETD2 protein that has been directed toward H3K36 through its association with RNAPII. In the latter situation, it is conceivable that all activity toward H3K36 can be inhibited by H3.3K36M while there is still SETD2 activity toward substrates other than H3K36 remaining, albeit that there is less total SETD2 activity available. This could explain why H3.3K36M does not affect protein content as strongly as SETD2 depletion despite a similar decrease in H3K36me3 levels.

If KDM4A and SETD2 regulate protein synthesis through a common pathway, it is plausible that SETD2 directly methylates a component of the ribosome, and that this methylation depends on an interaction between SETD2’s SRI domain and a component of the translation machinery. The SRI domain is positively charged at cellular pH (isoelectric point 8.97 for the SRI domain of human SETD2). Critical positively charged residues in the SRI domain (such as R2510) mediate the interaction with both the negatively charged RNAPII phosho-CTD (Li et al. 2005) as well as with the acidic C-terminal tail of alpha-tubulin (Kearns et al. 2020). Interestingly, a recent study on SETD2 interacting proteins identified the mRNA splicing regulating heterogeneous nuclear ribonucleoproteins (hnRNPs) as common SETD2 interactors (Bhattacharya et al. 2021). Among the other proteins identified as SETD2 interactors were also many ribosomal subunits. Although ribosomal proteins are common contaminants in co-IP experiments (Pardo and Choudhary 2012), it might be interesting to determine if SETD2 interacts with specific component of the ribosome via the SRI domain, and whether this interaction is relevant for the regulation of protein synthesis by SETD2.

In confirmation of our findings, a recent preprint study also found a negative role for SETD2 in translation regulation in clear cell renal cell carcinoma (ccRCC; Hapke et al. 2020). SETD2 inactivating mutations are frequently found in multiple types of cancer, including ccRCC (Dalgliesh et al. 2010; Duns et al. 2010; Gerlinger et al. 2012; Sato et al. 2013; Bihr et al. 2019), high-grade gliomas (Fontebasso et al. 2013), and leukemias (Zhang et al. 2012; Zhu et al. 2014; Mar et al. 2014). In addition, SETD2 is mutated at low frequency in many other types of cancers such as melanoma, and lung and colon adenocarcinoma (for review see Li et al. 2016; Fahey and Davis 2017; Chen et al. 2020). Perturbation of translation regulation is a common theme in cancer. Many tumor cells upregulate ribosome production and protein synthesis by overexpressing MYC, which promotes the expression of ribosome biogenesis genes (Muhar et al. 2018), and/or deregulating the RAS and PI3K signaling pathways (reviewed by Silvera et al. 2010; Robichaud et al. 2019). It is therefore tempting to speculate that SETD2 inactivation is another way for tumor cells to increase protein production. The tumor-suppressor function of SETD2 has so far been attributed to its role in DNA damage repair (Daugaard et al. 2012; Li et al. 2013; Carvalho et al. 2014; Pfister et al. 2014), transcription and mRNA processing (Simon et al. 2014; Grosso et al. 2015), and in genome stability (Park et al. 2016; Chiang et al. 2018). Our study warrants further investigation into the molecular mechanism of translation regulation by SETD2 as well as studies to determine if this function contributes to tumor development in SETD2 mutant or KDM4A overexpressing cancers.

## Materials and Methods

### Cell culture, knockdowns and overexpression

Human non-transformed retinal pigment epithelial cells transduced with the human telomerase gene (*hTERT*-RPE1; ATCC CRL-4000) were grown in DMEM/F12 (Gibco) supplemented with 10% fetal calf serum (FCS). TIG-3 cells (human diploid embryonic lung fibroblasts; Research Resource Identifier: CVCL_E939) and BJ cells (human diploid foreskin fibroblasts; ATCC CRL-2522) were previously transduced with *hTERT* and the murine ecotropic retrovirus receptor (Michaloglou et al. 2005). HEK293T, TIG-3 and BJ cells were grown in DMEM (Gibco) with 10% FCS. Cells were maintained at 37°C, 5% CO2 in a humidified incubator.

For microRNA (miRNA) based knockdown of SETD2, cells were lentivirally transduced with doxycycline (dox)-inducible artificial miRNAs in the miR-E backbone (Fellmann et al. 2013). *SETD2* targeting miRNA sequences were CCAGGACAGAAAGAAAGTTAGA (#1) and ACCGGAAGTTGTTTGAGCAAGA (#2). Non-targeting miRNA sequence was CAATGTACTGCGCGTGGAGACT. Knockdown was induced by treating cells with 1 μg/mL dox for 72h.

For CasRx based knockdown of SETD2, RPE1 cells were first transduced with a dox-inducible human codon-optimized CasRx construct (synthesized by Integrated DNA Technologies [IDT]) containing a blasticidin resistance gene. The CasRx protein sequence, including nuclear localization signal and hemagglutinin (HA) epitope tag, was identical as described in Konermann et al. (2018). After selection with 10 µg/mL blasticidin (Invivogen), a monoclonal cell line showing high CasRx expression after dox treatment was further transduced with SEDT2 gRNA#1 (AGATCCACAACAAAGACAGCCCA), SETD2 gRNA#2 (TTCACATTCTCATTGCACTCCAG) or a scrambled gRNA (TCACCAGAAGCGTACCATACTC) in a construct containing an enhanced green fluorescent protein (EGFP) marker. The SETD2 CasRx gRNAs were designed using the Cas13 guide design resource (Wessels et al. 2020). CasRx expression was induced by treating cells with 1 μg/mL dox for 72h.

For constitutive overexpression of SETD2, KDM4A, Rph1, and the yeast Set2 SRI domain, coding sequences were cloned into a lentiviral vector in which proteins are N-terminally tagged with tagRFP (Merzlyak et al. 2007) and expression is driven by the human core *EEF1A1* promoter. Coding sequences were followed by an internal ribosome entry site (IRES) sequence and a bleomycin/zeocin resistance gene. The coding sequence for truncated SETD2 (amino acids 504-2564) lacking part of the N-terminal unstructured domain was amplified from human RPE1-hTERT cDNA and made resistant to *SETD2* miRNA#1 and #2 by silent mutation of miRNA binding sites. Full-length *KDM4A* was amplified from human RPE1-hTERT cDNA. Full-length *RPH1* was amplified from genomic DNA from *Saccharomyces cerevisiae* strain BY4741. The SRI domain from *S. cerevisiae* Set2 (amino acids 619-733) was N-terminally tagged with an SV40 nuclear localization signal (NLS) or HIV Rev protein nuclear export signal (LPPLERLTL; NES) and codon optimized for expression in humans (synthesized by IDT). For expression of the human SETD2 SRI domain, cDNA derived sequences were N-terminally tagged with a destabilizing domain (DD; Banaszynski et al. 2006) that replaced the tagRFP followed by an HA tag and an SV40 NLS. To induce stabilization of DD tagged proteins, cells were treated with 0.5 μg/mL Shield-1 (Aobious) for 72h. For constitutive overexpression of H3.3 and H3.3K36A, codon optimized sequences with C-terminal HA epitope tags (synthesized by IDT) were cloned into the same lentiviral vector but without N-terminal tagRFP. Following transduction, cells were selected and maintained in medium with 100 μg/mL zeocin (Invivogen).

For dox inducible overexpression of H3.3, H3.3K36M and H3.3G34R, codon optimized coding sequences with a C-terminal HA epitope tag were synthesized by IDT and cloned into a pCW57.1 (Addgene plasmid #41393) derived lentiviral vector with a blasticidin resistance gene (replacing the original puromycin resistance gene). Following transduction, cells were selected with 10 µg/mL blasticidin (Invivogen) for 7 days. Overexpression was induced by treating cells with 1 μg/mL dox for 96h.

### Lentivirus production

Lentiviral transfer plasmids were co-transfected with pMD2G, pRSV-VSV and pMDL packaging plasmids in HEK293T cells using polyethyleminine (PEI) at a 1:3 DNA:PEI ratio. Supernatant was collected 48h and 72h post-transfection, passed through a 0.45 um filter and concentrated using an Amicon Ultra-15 centrifugal filter unit (UFC910024, Merck/Millipore), Ultracel-100 regenerated cellulose membrane.

### RNA isolation and RT-qPCR

RNA was isolated using the RNeasy Mini kit (Qiagen) with on-column DNAse I digestion. cDNA was synthesized using Superscript II Reverse Transcriptase (ThermoFisher) and random hexamers. For determining *SETD2* knockdown efficiency using the CasRx system, qPCR primers were designed around the gRNA target site. Primers for qPCR are listed in Table 1.

**Table 1.**
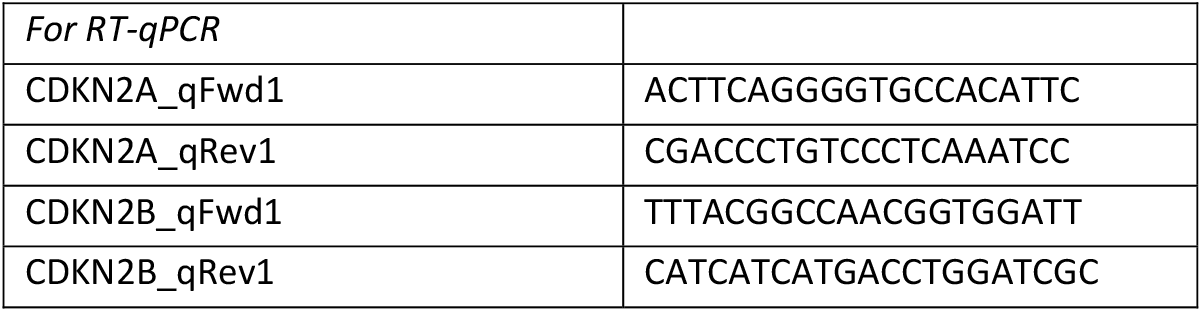
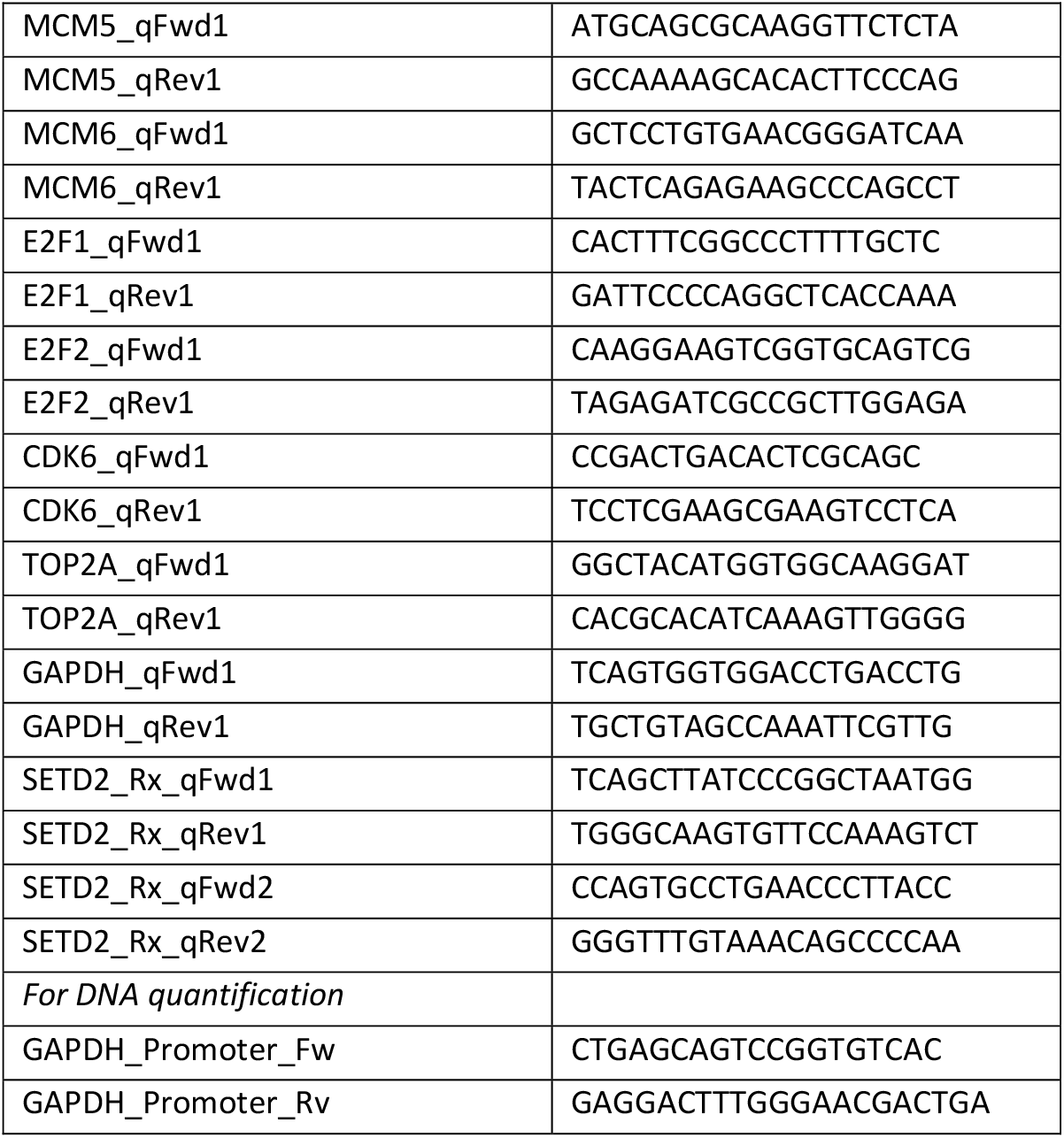
Primers used in this study.

### Co-immunoprecipitation

Plasmids encoding tagRFP or SRI_SETD2_ with N-terminal DD-HA-SV40 NLS fusions were transfected into HEK293T cells using Fugene HD at a 1:4 plasmid:FugeneHD ratio in OptiMEM. Cells were immediately treated with 0.5 μg/mL Shield-1 and harvested in IP lysis buffer (50 mM Tris-HCl pH 7.5, 150 mM NaCl, 5 mM EDTA, 0.5% IGEPAL, 1% Triton X-100) 48h after transfection. Cells were sonicated for 30 cycles at high setting (30s on, 30s off) using a Bioruptor Pico sonicator (Diagenode) and centrifuged at 13000 rpm for 10 min. Supernatant was used for immunoprecipitation with 5 μg anti-HA antibody overnight at 4°C. Next, immunocomplexes were precipitated with Protein G Dynabeads (ThermoFisher) for 4h at 4°C, washed three times with IP lysis buffer, and eluted with SDS loading buffer (50 mM Tris-HCl pH 6.8, 2% SDS, 10% glycerol, 0.1M dithiothreitol (DTT), 0.02% bromophenol blue). Samples were boiled, centrifuged and immunoprecipitated proteins were detected by Western blot.

### Western blot

Approximately 1×10^7^ cells were washed twice with phosphate-buffered saline (PBS). Proteins were isolated by adding SDS lysis buffer (50 mM Tris-HCl pH 6.8, 2% SDS, 10% glycerol) supplemented with protease inhibitor cocktail (PIC; Roche). DNA was sheared by sonication for 10 min at high settings (30 s on, 30 s off) using a Bioruptor Pico sonicator (Diagenode) to reduce sample viscosity. Protein concentration was determined with the DC protein assay (Bio-Rad) according to manufactures manual. Samples were supplemented with DTT (final 0.1M) and bromophenol blue (final 0.02%). Samples were boiled, centrifuged and 10 μg protein was separated on a NuPAGE 12% Bis-Tris protein gel (ThermoFisher) for histones and on a NuPAGE 4-12% Bis-Tris protein gel for non-histone proteins. Next, proteins were blotted on 0.2 μm (for histones) and 0.45 μm (for non-histone proteins) nitrocellulose membranes at 1 ampere for 90 min. Afterwards membranes were blocked for 30 min with 5% Nutrilon (Nutricia) in PBS and incubated overnight at 4°C with primary antibodies H3 (Abcam 1791), H4 (Merck Millipore 04-858), H4ac (Merck Millipore 06-866), H3K36me3 (Abcam 9050), H3K36me2 (gift from Dirk Schübeler), H3K9me3 (Abcam 8898), beta-actin (Abcam 6276), beta-actin (Santa Cruz sc-1616), alpha-tubulin (Sigma-Aldrich T5168), V5 (Invitrogen R960-25) and HA (Abcam 18181) in 2% Nutrilon in Tris-buffered saline-Tween (TBST). The next day membranes were washed four times with TBST before incubating the membrane with the appropriate Odyssey IRDye secondary antibody (LI-COR Biosciences) at 1:10000 dilution in 2% Nutrilon in TBST for 1h. Membranes were washed four times with TBST before scanning on a LI-COR Odyssey IR Imager (LI-COR Biosciences). Signals were quantified using Image Studio software (LI-COR).

To normalize protein lysates for genomic DNA concentration, aliquots of protein lysates with equal protein concentration were treated with proteinase K (ProtK) and RNAse A at 55°C for 30 min, followed by Proteinase K inactivation at 95°C for 10 min. DNA was ethanol precipitated, washed, dried and resuspended in 50 mM Tris-HCl pH 8. Relative genomic DNA concentrations were determined by qPCR using primers for the *GAPDH* promoter.

### ^35^S-methionine incorporation assay

Protein synthesis rates were measured as described previously (Faller et al. 2015). hTERT-RPE1 cells were incubated with DMEM methionine-free media (ThermoFisher Scientific #21013024) for 20 min, after which 30 µCi/ml ^35^S-methionine label (Hartmann Analytic) was added for 1 hour. After washing the samples with PBS, proteins were extracted with lysis buffer (50mM Tris-HCl pH 7.5, 150mM NaCl, 1% Tween-20, 0.5% NP-40, 1× protease inhibitor cocktail (Roche) and 1x phosphatase inhibitor cocktail (Sigma Aldrich) and precipitated onto filter paper (Whatmann) with 25% trichloroacetic acid and washed twice with 70% ethanol and twice with acetone. A liquid scintillation counter (Perkin Elmer) was used to measure scintillation and the activity was normalized by total protein content.

### Flow-cytometry

For cell cycle distribution analysis, hTERT-RPE1 cells were fixed for with 70% ethanol at 4°C for 30 min. Cells were treated with RNAse A and stained with propidium iodide (50 µg/ml). For image flow-cytometry, hTERT-RPE1 cells were detached from culture plates with accutase (Stemcell Technologies) and stained with CellTrace CFSE Cell Proliferation Kit C34554 (ThermoFisher) according to manufacturers’ protocol. 2D cell size was measured imaging flow-cytometry (ImageStream X Mark II).

### Statistical analysis

Statistical significance was calculated using a two-tailed, unpaired Student’s t-test.

## Declarations

### Availability of data and material

All processed data are within the paper and the Supplemental Material.

### Statistics

Statistical analyses were performed using GraphPad Prism 8. Data are presented as mean ± SD. Unless stated otherwise, the unpaired Student’s t-test with two-tailed distributions was used to calculate the p-value. A p-value < 0.05 was considered statistically significant.

### Author contributions

Conception and design: TMM and FvL

Acquisition of data: TMM, EMKM, JS, MM

Analysis and interpretation of data: TMM, EMKM, JS, MM, WJF, FvL

Supervision of experiments and analyses: WJF, FvL

Writing of the manuscript, TMM, EMKM, JS, MM, WJF, FvL

## Acknowledgements

We thank Marlize van Breugel for carefully reading the manuscript and for valuable suggestions, Dirk Schübeler (FMI) for providing the H3K36me2 antibody, Martijn van Baalen (NKI) for flow cytometry advice, and Erik Mul (Sanquin) for help with image flow-cytometry.

## Supplementary figures

**Supplementary Figure 1.**
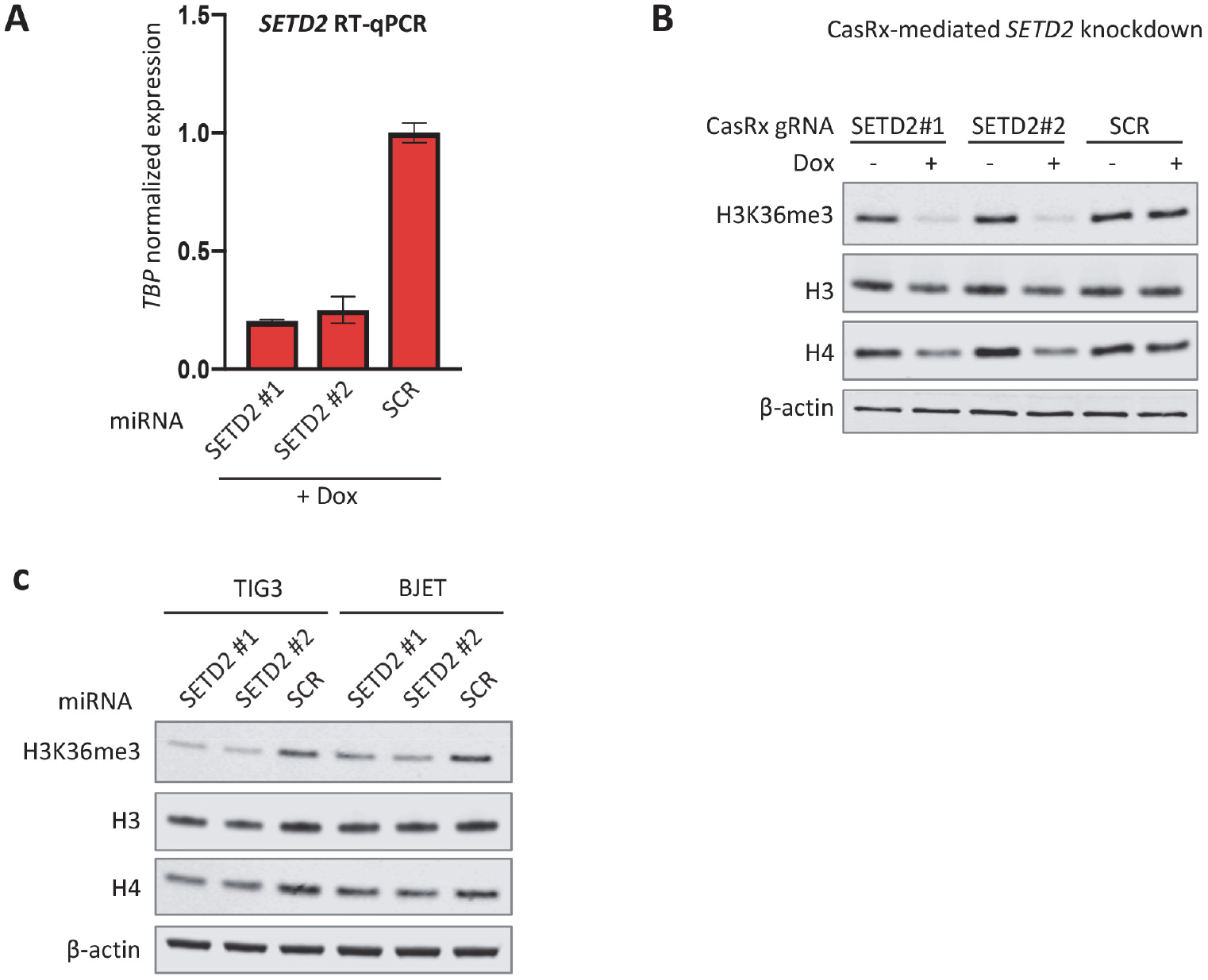
SETD2 knockdown using CasRx increased protein content in RPE1 cells and miRNA-based knockdown increases protein content in TIG3 and BJ cells. (**A**) RT-qPCR of *SETD2* following miRNA-based knockdown in RPE1 cells. (**B**) Western blot of RPE1 cells with doxycycline inducible expression of CasRx and stable expression of CasRx gRNAs targeting *SETD2* or a scrambled (SCR) gRNA. (**C**) Western blot of TIG3 and BJ cells with miRNA-based knockdown of SETD2.

## References

Banaszynski, Laura A., Ling-chun Chen, Lystranne A. Maynard-Smith, A. G. Lisa Ooi, and Thomas J. Wandless. 2006. “A Rapid, Reversible, and Tunable Method to Regulate Protein Function in Living Cells Using Synthetic Small Molecules.” Cell 126 (5): 995–1004. https://doi.org/10.1016/j.cell.2006.07.025.

Baubec, Tuncay, Daniele F. Colombo, Christiane Wirbelauer, Juliane Schmidt, Lukas Burger, Arnaud R. Krebs, Altuna Akalin, and Dirk Schübeler. 2015. “Genomic Profiling of DNA Methyltransferases Reveals a Role for DNMT3B in Genic Methylation.” Nature 520 (7546): 243–47. https://doi.org/10.1038/nature14176.

Behjati, Sam, Patrick S. Tarpey, Nadège Presneau, Susanne Scheipl, Nischalan Pillay, Peter Van Loo, David C. Wedge, et al. 2013. “Distinct H3F3A and H3F3B Driver Variants Define Chondroblastoma and Giant Cell Tumour of Bone.” Nature Genetics 45 (12). https://doi.org/10.1038/ng.2814.

Bertomeu, Thierry, Jasmin Coulombe-Huntington, Andrew Chatr-aryamontri, Karine G. Bourdages, Etienne Coyaud, Brian Raught, Yu Xia, and Mike Tyers. 2018. “A High-Resolution Genome-Wide CRISPR/Cas9 Viability Screen Reveals Structural Features and Contextual Diversity of the Human Cell-Essential Proteome.” Molecular and Cellular Biology 38 (1). https://doi.org/10.1128/MCB.00302-17.

Bhattacharya, Saikat, Michaella J. Levy, Ning Zhang, Hua Li, Laurence Florens, Michael P. Washburn, and Jerry L. Workman. 2021. “The Methyltransferase SETD2 Couples Transcription and Splicing by Engaging MRNA Processing Factors through Its SHI Domain.” Nature Communications 12 (1): 1443. https://doi.org/10.1038/s41467-021-21663-w.

Bihr, Svenja, Riuko Ohashi, Ariane L. Moore, Jan H. Rüschoff, Christian Beisel, Thomas Hermanns, Axel Mischo, et al. 2019. “Expression and Mutation Patterns of PBRM1, BAP1 and SETD2 Mirror Specific Evolutionary Subtypes in Clear Cell Renal Cell Carcinoma.” Neoplasia (New York, N.Y.) 21 (2): 247–56. https://doi.org/10.1016/j.neo.2018.12.006.

Blomen, Vincent A., Peter Májek, Lucas T. Jae, Johannes W. Bigenzahn, Joppe Nieuwenhuis, Jacqueline Staring, Roberto Sacco, et al. 2015. “Gene Essentiality and Synthetic Lethality in Haploid Human Cells.” Science 350 (6264): 1092–96. https://doi.org/10.1126/science.aac7557.

Bondarenko, Vladimir A., Louise M. Steele, Andrea Újvári, Daria A. Gaykalova, Olga I. Kulaeva, Yury S. Polikanov, Donal S. Luse, and Vasily M. Studitsky. 2006. “Nucleosomes Can Form a Polar Barrier to Transcript Elongation by RNA Polymerase II.” Molecular Cell 24 (3): 469–79. https://doi.org/10.1016/j.molcel.2006.09.009.

Carrozza, Michael J., Bing Li, Laurence Florens, Tamaki Suganuma, Selene K. Swanson, Kenneth K. Lee, Wei-Jong Shia, et al. 2005. “Histone H3 Methylation by Set2 Directs Deacetylation of Coding Regions by Rpd3S to Suppress Spurious Intragenic Transcription.” Cell 123 (4): 581–92. https://doi.org/10.1016/j.cell.2005.10.023.

Carvalho Sílvia, Ana Cláudia Raposo, Filipa Batalha Martins, Ana Rita Grosso, Sreerama Chaitanya Sridhara, José Rino, Maria Carmo-Fonseca, and Sérgio Fernandes de Almeida. 2013. “Histone Methyltransferase SETD2 Coordinates FACT Recruitment with Nucleosome Dynamics during Transcription.” Nucleic Acids Research 41 (5): 2881–93. https://doi.org/10.1093/nar/gks1472.

Carvalho Sílvia, Alexandra C. Vítor, Sreerama C. Sridhara, Filipa B. Martins, Ana C. Raposo, Joana MP Desterro, João Ferreira, and Sérgio F. de Almeida. 2014. “SETD2 Is Required for DNA Double-Strand Break Repair and Activation of the P53-Mediated Checkpoint.” ELife 3. https://doi.org/10.7554/eLife.02482.

Chen, Kun, Juan Liu, Shuxun Liu, Meng Xia, Xiaomin Zhang, Dan Han, Yingming Jiang, Chunmei Wang, and Xuetao Cao. 2017. “Methyltransferase SETD2-Mediated Methylation of STAT1 Is Critical for Interferon Antiviral Activity.” Cell 170 (3): 492-506.e14. https://doi.org/10.1016/j.cell.2017.06.042.

Chen, Rui, Wei-qing Zhao, Cheng Fang, Xin Yang, and Mei Ji. 2020. “Histone Methyltransferase SETD2: A Potential Tumor Suppressor in Solid Cancers.” Journal of Cancer 11 (11): 3349–56. https://doi.org/10.7150/jca.38391.

Chen, Zhijie, Ronen Gabizon, Aidan I Brown, Antony Lee, Aixin Song, César Díaz-Celis, Craig D Kaplan, Elena F Koslover, Tingting Yao, and Carlos Bustamante. 2019. “High-Resolution and High-Accuracy Topographic and Transcriptional Maps of the Nucleosome Barrier.” Edited by Taekjip Ha and Jessica K Tyler. ELife 8 (July): e48281. https://doi.org/10.7554/eLife.48281.

Chiang, Yun-Chen, In-Young Park, Esteban A. Terzo, Durga Nand Tripathi, Frank M. Mason, Catherine C. Fahey, Menuka Karki, et al. 2018. “SETD2 Haploinsufficiency for Microtubule Methylation Is an Early Driver of Genomic Instability in Renal Cell Carcinoma.” Cancer Research 78 (12): 3135–46. https://doi.org/10.1158/0008-5472.CAN-17-3460.

Conlon, Ian, and Martin Raff. 2003. “Differences in the Way a Mammalian Cell and Yeast Cells Coordinate Cell Growth and Cell-Cycle Progression.” Journal of Biology 2 (1): 7. https://doi.org/10.1186/1475-4924-2-7.

Dalgliesh, Gillian L., Kyle Furge, Chris Greenman, Lina Chen, Graham Bignell, Adam Butler, Helen Davies, et al. 2010. “Systematic Sequencing of Renal Carcinoma Reveals Inactivation of Histone Modifying Genes.” Nature 463 (7279): 360–63. https://doi.org/10.1038/nature08672.

Daugaard, Mads, Annika Baude, Kasper Fugger, Lou Klitgaard Povlsen, Halfdan Beck, Claus Storgaard Sørensen, Nikolaj H. T. Petersen, et al. 2012. “LEDGF (P75) Promotes DNA-End Resection and Homologous Recombination.” Nature Structural & Molecular Biology 19 (8): 803–10. https://doi.org/10.1038/nsmb.2314.

Duns, Gerben, Eva van den Berg, Inge van Duivenbode, Jan Osinga, Harry Hollema, Robert M. W. Hofstra, and Klaas Kok. 2010. “Histone Methyltransferase Gene SETD2 Is a Novel Tumor Suppressor Gene in Clear Cell Renal Cell Carcinoma.” Cancer Research 70 (11): 4287–91. https://doi.org/10.1158/0008-5472.CAN-10-0120.

Edmunds, John W., Louis C. Mahadevan, and Alison L. Clayton. 2008. “Dynamic Histone H3 Methylation during Gene Induction: HYPB/Setd2 Mediates All H3K36 Trimethylation.” The EMBO Journal 27 (2): 406–20. https://doi.org/10.1038/sj.emboj.7601967.

Fahey, Catherine C., and Ian J. Davis. 2017. “SETting the Stage for Cancer Development: SETD2 and the Consequences of Lost Methylation.” Cold Spring Harbor Perspectives in Medicine 7 (5): a026468. https://doi.org/10.1101/cshperspect.a026468.

Faller, William J., Thomas J. Jackson, John Rp Knight, Rachel A. Ridgway, Thomas Jamieson, Saadia A. Karim, Carolyn Jones, et al. 2015. “MTORC1-Mediated Translational Elongation Limits Intestinal Tumour Initiation and Growth.” Nature 517 (7535): 497–500. https://doi.org/10.1038/nature13896.

Fang, Dong, Haiyun Gan, Jeong-Heon Lee, Jing Han, Zhiquan Wang, Scott M. Riester, Long Jin, et al. 2016. “The Histone H3.3K36M Mutation Reprograms the Epigenome of Chondroblastomas.” Science (New York, N.Y.) 352 (6291): 1344. https://doi.org/10.1126/science.aae0065.

Fang, Jun, Yaping Huang, Guogen Mao, Shuang Yang, Gadi Rennert, Liya Gu, Haitao Li, and Guo-Min Li. 2018. “Cancer-Driving H3G34V/R/D Mutations Block H3K36 Methylation and H3K36me3–MutSα Interaction.” Proceedings of the National Academy of Sciences of the United States of America 115 (38): 9598. https://doi.org/10.1073/pnas.1806355115.

Farnung, Lucas, Moritz Ochmann, Maik Engeholm, and Patrick Cramer. 2021. “Structural Basis of Nucleosome Transcription Mediated by Chd1 and FACT.” Nature Structural & Molecular Biology 28 (4): 382–87. https://doi.org/10.1038/s41594-021-00578-6.

Fellmann, Christof, Thomas Hoffmann, Vaishali Sridhar, Barbara Hopfgartner, Matthias Muhar, Mareike Roth, Dan Yu Lai, et al. 2013. “An Optimized MicroRNA Backbone for Effective Single- Copy RNAi.” Cell Reports 5 (6): 1704–13. https://doi.org/10.1016/j.celrep.2013.11.020.

Fontebasso, Adam M., Jeremy Schwartzentruber, Dong-Anh Khuong-Quang, Xiao-Yang Liu, Dominik Sturm, Andrey Korshunov, David T. W. Jones, et al. 2013. “Mutations in SETD2 and Genes Affecting Histone H3K36 Methylation Target Hemispheric High-Grade Gliomas.” Acta Neuropathologica 125 (5): 659–69. https://doi.org/10.1007/s00401-013-1095-8.

Gerlinger, Marco, Andrew J. Rowan, Stuart Horswell, James Larkin, David Endesfelder, Eva Gronroos, Pierre Martinez, et al. 2012. “Intratumor Heterogeneity and Branched Evolution Revealed by Multiregion Sequencing.” New England Journal of Medicine 366 (10): 883–92. https://doi.org/10.1056/NEJMoa1113205.

Gopalakrishnan, Rajaraman, Sharon K Marr, Robert E Kingston, and Fred Winston. 2019. “A Conserved Genetic Interaction between Spt6 and Set2 Regulates H3K36 Methylation.” Nucleic Acids Research 47 (8): 3888–3903. https://doi.org/10.1093/nar/gkz119.

Goranov, Alexi I., Michael Cook, Marketa Ricicova, Giora Ben-Ari, Christian Gonzalez, Carl Hansen, Mike Tyers, and Angelika Amon. 2009. “The Rate of Cell Growth Is Governed by Cell Cycle Stage.” Genes & Development 23 (12): 1408–22. https://doi.org/10.1101/gad.1777309.

Grosso, Ana R., Ana P. Leite, Sílvia Carvalho, Mafalda R. Matos, Filipa B. Martins Alexandra, C. Vítor, Joana M. P. Desterro, Maria Carmo-Fonseca, and Sérgio F. de Almeida. 2015. “Pervasive Transcription Read-through Promotes Aberrant Expression of Oncogenes and RNA Chimeras in Renal Carcinoma.” ELife 4 (November). https://doi.org/10.7554/eLife.09214.

Guo, Rui, Lijuan Zheng, Juw Won Park, Ruitu Lv, Hao Chen, Fangfang Jiao, Wenqi Xu, et al. 2014. “BS69/ZMYND11 Reads and Connects Histone H3.3 Lysine 36 Trimethylation Decorated Chromatin to Regulated Pre-MRNA Processing.” Molecular Cell 56 (2): 298–310. https://doi.org/10.1016/j.molcel.2014.08.022.

Hacker, Kathryn E., Catherine C. Fahey, Stephen A. Shinsky, Yun-Chen J. Chiang, Julia V. DiFiore, Deepak Kumar Jha, Andy H. Vo, et al. 2016. “Structure/Function Analysis of Recurrent Mutations in SETD2 Protein Reveals a Critical and Conserved Role for a SET Domain Residue in Maintaining Protein Stability and Histone H3 Lys-36 Trimethylation.” The Journal of Biological Chemistry 291 (40): 21283–95. https://doi.org/10.1074/jbc.M116.739375.

Hapke, Robert, Lindsay Venton, Kristie Lindsay Rose, Quanhu Sheng, Anupama Reddy, Angela Jones, W. Kimryn Rathmell, and Scott Haake. 2020. “SETD2 Regulates the Methylation of Translation Elongation Factor EEF1A1 in Clear Cell Renal Cell Carcinoma.” Preprint. Cancer Biology. https://doi.org/10.1101/2020.10.26.354902.

Hsieh, Fu-Kai, Olga I. Kulaeva, Smita S. Patel, Pamela N. Dyer, Karolin Luger, Danny Reinberg, and Vasily M. Studitsky. 2013. “Histone Chaperone FACT Action during Transcription through Chromatin by RNA Polymerase II.” Proceedings of the National Academy of Sciences 110 (19): 7654–59. https://doi.org/10.1073/pnas.1222198110.

Huang, Yaping, Liya Gu, and Guo-Min Li. 2018. “H3K36me3-Mediated Mismatch Repair Preferentially Protects Actively Transcribed Genes from Mutation.” Journal of Biological Chemistry 293 (20): 7811–23. https://doi.org/10.1074/jbc.RA118.002839.

Hyun, Kwangbeom, Jongcheol Jeon, Kihyun Park, and Jaehoon Kim. 2017. “Writing, Erasing and Reading Histone Lysine Methylations.” Experimental & Molecular Medicine 49 (4): e324–e324. https://doi.org/10.1038/emm.2017.11.

Jamai, Adil, Andrea Puglisi, and Michel Strubin. 2009. “Histone Chaperone Spt16 Promotes Redeposition of the Original H3-H4 Histones Evicted by Elongating RNA Polymerase.” Molecular Cell 35 (3): 377–83. https://doi.org/10.1016/j.molcel.2009.07.001.

Jeronimo, Célia, Christian Poitras, and François Robert. 2019. “Histone Recycling by FACT and Spt6 during Transcription Prevents the Scrambling of Histone Modifications.” Cell Reports 28 (5): 1206-1218.e8. https://doi.org/10.1016/j.celrep.2019.06.097.

Joshi, Amita A., and Kevin Struhl. 2005. “Eaf3 Chromodomain Interaction with Methylated H3-K36 Links Histone Deacetylation to Pol II Elongation.” Molecular Cell 20 (6): 971–78. https://doi.org/10.1016/j.molcel.2005.11.021.

Kearns, Sarah, Frank M. Mason, W. Kimryn Rathmell, In Young Park, Cheryl Walker, Kristen Verhey, and Michael A. Cianfrocco. 2020. “Molecular Determinants for α-Tubulin Methylation by SETD2.” Preprint. Biochemistry. https://doi.org/10.1101/2020.10.21.349365.

Keogh, Michael-Christopher, Siavash K. Kurdistani, Stephanie A. Morris, Seong Hoon Ahn, Vladimir Podolny, Sean R. Collins, Maya Schuldiner, et al. 2005. “Cotranscriptional Set2 Methylation of Histone H3 Lysine 36 Recruits a Repressive Rpd3 Complex.” Cell 123 (4): 593–605. https://doi.org/10.1016/j.cell.2005.10.025.

Kim, TaeSoo, and Stephen Buratowski. 2007. “Two Saccharomyces Cerevisiae JmjC Domain Proteins Demethylate Histone H3 Lys36 in Transcribed Regions to Promote Elongation.” Journal of Biological Chemistry 282 (29): 20827–35. https://doi.org/10.1074/jbc.M703034200.

Kizer, Kelby O., Hemali P. Phatnani, Yoichiro Shibata, Hana Hall, Arno L. Greenleaf, and Brian D. Strahl. 2005. “A Novel Domain in Set2 Mediates RNA Polymerase II Interaction and Couples Histone H3 K36 Methylation with Transcript Elongation.” Molecular and Cellular Biology 25 (8): 3305–16. https://doi.org/10.1128/MCB.25.8.3305-3316.2005.

Klose, Robert J., Kathryn E. Gardner, Gaoyang Liang, Hediye Erdjument-Bromage, Paul Tempst, and Yi Zhang. 2007. “Demethylation of Histone H3K36 and H3K9 by Rph1: A Vestige of an H3K9 Methylation System in Saccharomyces Cerevisiae?” Molecular and Cellular Biology 27 (11): 3951–61. https://doi.org/10.1128/MCB.02180-06.

Konermann, Silvana, Peter Lotfy, Nicholas J. Brideau, Jennifer Oki, Maxim N. Shokhirev, and Patrick D. Hsu. 2018. “Transcriptome Engineering with RNA-Targeting Type VI-D CRISPR Effectors.” Cell 173 (3): 665-676.e14. https://doi.org/10.1016/j.cell.2018.02.033.

LeRoy, Gary, Peter A. DiMaggio, Eric Y. Chan, Barry M. Zee, M. Andres Blanco, Barbara Bryant, Ian Z. Flaniken, et al. 2013. “A Quantitative Atlas of Histone Modification Signatures from Human Cancer Cells.” Epigenetics & Chromatin 6 (1): 20. https://doi.org/10.1186/1756-8935-6-20.

Leung, Calvin S., Stephen M. Douglass, Marco Morselli, Matthew B. Obusan, Marat S. Pavlyukov, Matteo Pellegrini, and Tracy L. Johnson. 2019. “H3K36 Methylation and the Chromodomain Protein Eaf3 Are Required for Proper Cotranscriptional Spliceosome Assembly.” Cell Reports 27 (13): 3760-3769.e4. https://doi.org/10.1016/j.celrep.2019.05.100.

Lewis, Peter W., Manuel M. Müller, Matthew S. Koletsky, Francisco Cordero, Shu Lin, Laura A. Banaszynski, Benjamin A. Garcia, Tom W. Muir, Oren J. Becher, and C. David Allis. 2013. “Inhibition of PRC2 Activity by a Gain-of-Function H3 Mutation Found in Pediatric Glioblastoma.” Science 340 (6134): 857–61. https://doi.org/10.1126/science.1232245.

Li, Feng, Guogen Mao, Dan Tong, Jian Huang, Liya Gu, Wei Yang, and Guo-Min Li. 2013. “The Histone Mark H3K36me3 Regulates Human DNA Mismatch Repair through Its Interaction with MutSα.” Cell 153 (3): 590–600. https://doi.org/10.1016/j.cell.2013.03.025.

Li, Jie, Jeong Hyun Ahn, and Gang Greg Wang. 2019. “Understanding Histone H3 Lysine 36 Methylation and Its Deregulation in Disease.” Cellular and Molecular Life Sciences 76 (15): 2899–2916. https://doi.org/10.1007/s00018-019-03144-y.

Li, Jun, Gerben Duns, Helga Westers, Rolf Sijmons, Anke van den Berg, and Klaas Kok. 2016. “SETD2: An Epigenetic Modifier with Tumor Suppressor Functionality.” Oncotarget 7 (31): 50719–34. https://doi.org/10.18632/oncotarget.9368.

Li, Ming, Hemali P. Phatnani, Ziqiang Guan, Harvey Sage, Arno L. Greenleaf, and Pei Zhou. 2005. “Solution Structure of the Set2–Rpb1 Interacting Domain of Human Set2 and Its Interaction with the Hyperphosphorylated C-Terminal Domain of Rpb1.” Proceedings of the National Academy of Sciences 102 (49): 17636–41. https://doi.org/10.1073/pnas.0506350102.

Lickwar, Colin R., Bhargavi Rao, Andrey A. Shabalin, Andrew B. Nobel, Brian D. Strahl, and Jason D. Lieb. 2009. “The Set2/Rpd3S Pathway Suppresses Cryptic Transcription without Regard to Gene Length or Transcription Frequency.” PLOS ONE 4 (3): e4886. https://doi.org/10.1371/journal.pone.0004886.

Lu, Chao, Siddhant U. Jain, Dominik Hoelper, Denise Bechet, Rosalynn C. Molden, Leili Ran, Devan Murphy, et al. 2016. “Histone H3K36 Mutations Promote Sarcomagenesis through Altered Histone Methylation Landscape.” Science (New York, N.Y.) 352 (6287): 844–49. https://doi.org/10.1126/science.aac7272.

Lu, Mingdong, Bin Zhao, Mengshan Liu, L. Wu, Yingying Li, Yingna Zhai, and Xian Shen. 2021. “Pan-Cancer Analysis of SETD2 Mutation and Its Association with the Efficacy of Immunotherapy.” Npj Precision Oncology 5 (1): 1–6. https://doi.org/10.1038/s41698-021-00193-0.

Luco, Reini F., Qun Pan, Kaoru Tominaga, Benjamin J. Blencowe, Olivia M. Pereira-Smith, and Tom Misteli. 2010. “Regulation of Alternative Splicing by Histone Modifications.” Science (New York, N.Y.) 327 (5968): 996–1000. https://doi.org/10.1126/science.1184208.

Mar, Brenton G., Lars B. Bullinger, Kathleen M. McLean, Peter V. Grauman, Marian H. Harris, Kristen Stevenson, Donna S. Neuberg, et al. 2014. “Mutations in Epigenetic Regulators Including SETD2 Are Gained during Relapse in Paediatric Acute Lymphoblastic Leukaemia.” Nature Communications 5 (March): 3469. https://doi.org/10.1038/ncomms4469.

Mar, Brenton G., S. Haihua Chu, Josephine D. Kahn, Andrei V. Krivtsov, Richard Koche, Cecilia A. Castellano, Jacob L. Kotlier, et al. 2017. “SETD2 Alterations Impair DNA Damage Recognition and Lead to Resistance to Chemotherapy in Leukemia.” Blood 130 (24): 2631. https://doi.org/10.1182/blood-2017-03-775569.

Maze, Ian, Wendy Wenderski, Kyung-Min Noh, Rosemary C. Bagot, Nikos Tzavaras, Immanuel Purushothaman, Simon J. Elsässer, et al. 2015. “Critical Role of Histone Turnover in Neuronal Transcription and Plasticity.” Neuron 87 (1): 77–94. https://doi.org/10.1016/j.neuron.2015.06.014.

McDaniel, Stephen L., and Brian D. Strahl. 2017. “Shaping the Cellular Landscape with Set2/SETD2 Methylation.” Cellular and Molecular Life Sciences 74 (18): 3317–34. https://doi.org/10.1007/s00018-017-2517-x.

Meers, Michael P, Telmo Henriques, Christopher A Lavender, Daniel J McKay, Brian D Strahl, Robert J Duronio, Karen Adelman, and A Gregory Matera. 2017. “Histone Gene Replacement Reveals a Post-Transcriptional Role for H3K36 in Maintaining Metazoan Transcriptome Fidelity.” Edited by Elisa Izaurralde. ELife 6 (March): e23249. https://doi.org/10.7554/eLife.23249.

Michaloglou, Chrysiis, Liesbeth C. W. Vredeveld, Maria S. Soengas, Christophe Denoyelle, Thomas Kuilman, Chantal M. A. M. van der Horst, Donné M. Majoor, Jerry W. Shay, Wolter J. Mooi, and Daniel S. Peeper. 2005. “BRAFE600-Associated Senescence-like Cell Cycle Arrest of Human Naevi.” Nature 436 (7051): 720–24. https://doi.org/10.1038/nature03890.

Molenaar, Thom M., Marc Pagès-Gallego, Vanessa Meyn, and Fred van Leeuwen. 2020. “Application of Recombination -Induced Tag Exchange (RITE) to Study Histone Dynamics in Human Cells.” Epigenetics 15 (9): 901–13. https://doi.org/10.1080/15592294.2020.1741777.

Muhar, Matthias, Anja Ebert, Tobias Neumann, Christian Umkehrer, Julian Jude, Corinna Wieshofer, Philipp Rescheneder, et al. 2018. “SLAM-Seq Defines Direct Gene-Regulatory Functions of the BRD4-MYC Axis.” Science (New York, N.Y.) 360 (6390): 800–805. https://doi.org/10.1126/science.aao2793.

Neri, Francesco, Stefania Rapelli, Anna Krepelova, Danny Incarnato, Caterina Parlato, Giulia Basile, Mara Maldotti, Francesca Anselmi, and Salvatore Oliviero. 2017. “Intragenic DNA Methylation Prevents Spurious Transcription Initiation.” Nature 543 (7643): 72–77. https://doi.org/10.1038/nature21373.

Pardo, Mercedes, and Jyoti S. Choudhary. 2012. “Assignment of Protein Interactions from Affinity Purification/Mass Spectrometry Data.” Journal of Proteome Research 11 (3): 1462–74. https://doi.org/10.1021/pr2011632.

Park, Young, Reid T. Powell, Durga Nand Tripathi, Ruhee Dere, Thai H. Ho, T. Lynne Blasius, Yun-Chen Chiang, et al. 2016. “Dual Chromatin and Cytoskeletal Remodeling by SETD2.” Cell 166 (4): 950. https://doi.org/10.1016/j.cell.2016.07.005.

Petesch, Steven J., and John T. Lis. 2012. “Overcoming the Nucleosome Barrier During Transcript Elongation.” Trends in Genetics : TIG 28 (6): 285–94. https://doi.org/10.1016/j.tig.2012.02.005.

Pfister, Sophia X., Sara Ahrabi, Lykourgos-Panagiotis Zalmas, Sovan Sarkar, François Aymard, Csanád Z. Bachrati, Thomas Helleday, et al. 2014. “SETD2-Dependent Histone H3K36 Trimethylation Is Required for Homologous Recombination Repair and Genome Stability.” Cell Reports 7 (6): 2006–18. https://doi.org/10.1016/j.celrep.2014.05.026.

Ponnaluri, V. K. Chaithanya, Divya Teja Vavilala, Sandeep Putty, William G. Gutheil, and Mridul Mukherji. 2009. “Identification of Non-Histone Substrates for JMJD2A-C Histone Demethylases.” Biochemical and Biophysical Research Communications 390 (2): 280–84. https://doi.org/10.1016/j.bbrc.2009.09.107.

Radman-Livaja, Marta, Tiffani K. Quan, Lourdes Valenzuela, Jennifer A. Armstrong, Tibor van Welsem, TaeSoo Kim, Laura J. Lee, et al. 2012. “A Key Role for Chd1 in Histone H3 Dynamics at the 3′ Ends of Long Genes in Yeast.” PLOS Genetics 8 (7): e1002811. https://doi.org/10.1371/journal.pgen.1002811.

Ray-Gallet, Dominique, Jean-Pierre Quivy, Christine Scamps, Emmanuelle M. -D Martini, Marc Lipinski, and Geneviève Almouzni. 2002. “HIRA Is Critical for a Nucleosome Assembly Pathway Independent of DNA Synthesis.” Molecular Cell 9 (5): 1091–1100. https://doi.org/10.1016/S1097-2765(02)00526-9.

Rebehmed, Joseph, Patrick Revy, Guilhem Faure, Jean-Pierre de Villartay, and Isabelle Callebaut. 2014. “Expanding the SRI Domain Family: A Common Scaffold for Binding the Phosphorylated C- Terminal Domain of RNA Polymerase II.” FEBS Letters 588 (23): 4431–37. https://doi.org/10.1016/j.febslet.2014.10.014.

Robichaud, Nathaniel, Nahum Sonenberg, Davide Ruggero, and Robert J. Schneider. 2019. “Translational Control in Cancer.” Cold Spring Harbor Perspectives in Biology 11 (7): a032896. https://doi.org/10.1101/cshperspect.a032896.

Sankaran, Saumya M., and Or Gozani. 2017. “Characterization of H3.3K36M as a Tool to Study H3K36 Methylation in Cancer Cells.” Epigenetics 12 (11): 917. https://doi.org/10.1080/15592294.2017.1377870.

Sato, Yusuke, Tetsuichi Yoshizato, Yuichi Shiraishi, Shigekatsu Maekawa, Yusuke Okuno, Takumi Kamura, Teppei Shimamura, et al. 2013. “Integrated Molecular Analysis of Clear-Cell Renal Cell Carcinoma.” Nature Genetics 45 (8): 860–67. https://doi.org/10.1038/ng.2699.

Schwartzentruber, Jeremy, Andrey Korshunov, Xiao-Yang Liu, David T. W. Jones, Elke Pfaff, Karine Jacob, Dominik Sturm, et al. 2012. “Driver Mutations in Histone H3.3 and Chromatin Remodelling Genes in Paediatric Glioblastoma.” Nature 482 (7384): 226–31. https://doi.org/10.1038/nature10833.

Seervai, Riyad N. H., Rahul K. Jangid, Menuka Karki, Durga Nand Tripathi, Sung Yun Jung, Sarah E. Kearns, Kristen J. Verhey, et al. 2020. “The Huntingtin-Interacting Protein SETD2/HYPB Is an Actin Lysine Methyltransferase.” Science Advances 6 (40). https://doi.org/10.1126/sciadv.abb7854.

Shi, Leilei, Jiejun Shi, Xiaobing Shi, Wei Li, and Hong Wen. 2018. “Histone H3.3 G34 Mutations Alter Histone H3K36 and H3K27 Methylation In Cis.” Journal of Molecular Biology 430 (11): 1562–65. https://doi.org/10.1016/j.jmb.2018.04.014.

Silvera, Deborah, Silvia C. Formenti, and Robert J. Schneider. 2010. “Translational Control in Cancer.” Nature Reviews Cancer 10 (4): 254–66. https://doi.org/10.1038/nrc2824.

Simon, Jeremy M., Kathryn E. Hacker, Darshan Singh, A. Rose Brannon, Joel S. Parker, Matthew Weiser, Thai H. Ho, et al. 2014. “Variation in Chromatin Accessibility in Human Kidney Cancer Links H3K36 Methyltransferase Loss with Widespread RNA Processing Defects.” Genome Research 24 (2): 241–50. https://doi.org/10.1101/gr.158253.113.

Smolle, Michaela, Swaminathan Venkatesh, Madelaine M. Gogol, Hua Li, Ying Zhang, Laurence Florens, Michael P. Washburn, and Jerry L. Workman. 2012. “Chromatin Remodelers Isw1 and Chd1 Maintain Chromatin Structure during Transcription by Preventing Histone Exchange.” Nature Structural & Molecular Biology 19 (9): 884–92. https://doi.org/10.1038/nsmb.2312.

Sorenson, Matthew R., Deepak K. Jha, Stefanie A. Ucles, Danielle M. Flood, Brian D. Strahl, Scott W. Stevens, and Tracy L. Kress. 2016. “Histone H3K36 Methylation Regulates Pre-MRNA Splicing in Saccharomyces Cerevisiae.” RNA Biology 13 (4): 412–26. https://doi.org/10.1080/15476286.2016.1144009.

Strahl, Brian D., Patrick A. Grant, Scott D. Briggs, Zu-Wen Sun, James R. Bone, Jennifer A. Caldwell, Sahana Mollah, et al. 2002. “Set2 Is a Nucleosomal Histone H3-Selective Methyltransferase That Mediates Transcriptional Repression.” Molecular and Cellular Biology 22 (5): 1298–1306. https://doi.org/10.1128/mcb.22.5.1298-1306.2002.

Studitsky, Vasily M., Ekaterina V. Nizovtseva, Alexey K. Shaytan, and Donal S. Luse. 2016. “Nucleosomal Barrier to Transcription: Structural Determinants and Changes in Chromatin Structure.” Biochemistry & Molecular Biology Journal 2 (2): 8. https://doi.org/10.21767/2471-8084.100017.

Sun, Xiao-Jian, Ju Wei, Xin-Yan Wu, Ming Hu, Lan Wang, Hai-Hong Wang, Qing-Hua Zhang, Sai-Juan Chen, Qiu-Hua Huang, and Zhu Chen. 2005. “Identification and Characterization of a Novel Human Histone H3 Lysine 36-Specific Methyltransferase.” The Journal of Biological Chemistry 280 (42): 35261–71. https://doi.org/10.1074/jbc.M504012200.

Tagami, Hideaki, Dominique Ray-Gallet, Geneviève Almouzni, and Yoshihiro Nakatani. 2004. “Histone H3.1 and H3.3 Complexes Mediate Nucleosome Assembly Pathways Dependent or Independent of DNA Synthesis.” Cell 116 (1): 51–61. https://doi.org/10.1016/S0092-8674(03)01064-X.

Tvardovskiy, Andrey, Veit Schwämmle, Stefan J. Kempf, Adelina Rogowska-Wrzesinska, and Ole N. Jensen. 2017. “Accumulation of Histone Variant H3.3 with Age Is Associated with Profound Changes in the Histone Methylation Landscape.” Nucleic Acids Research 45 (16): 9272–89. https://doi.org/10.1093/nar/gkx696.

Van Rechem, Capucine, Joshua C. Black, Myriam Boukhali, Martin J. Aryee, Susanne Gräslund, Wilhelm Haas, Cyril H. Benes, and Johnathan R. Whetstine. 2015. “Lysine Demethylase KDM4A Associates with Translation Machinery and Regulates Protein Synthesis.” Cancer Discovery 5 (3): 255–63. https://doi.org/10.1158/2159-8290.CD-14-1326.

Venkatesh, Swaminathan, Michaela Smolle, Hua Li, Madelaine M. Gogol, Malika Saint, Shambhu Kumar, Krishnamurthy Natarajan, and Jerry L. Workman. 2012. “Set2 Methylation of Histone H3 Lysine 36 Suppresses Histone Exchange on Transcribed Genes.” Nature 489 (7416): 452–55. https://doi.org/10.1038/nature11326.

Wagner, Eric J., and Phillip B. Carpenter. 2012. “Understanding the Language of Lys36 Methylation at Histone H3.” Nature Reviews. Molecular Cell Biology 13 (2): 115–26. https://doi.org/10.1038/nrm3274.

Wang, Tim, Kıvanç Birsoy, Nicholas W. Hughes, Kevin M. Krupczak, Yorick Post, Jenny J. Wei, Eric S. Lander, and David M. Sabatini. 2015. “Identification and Characterization of Essential Genes in the Human Genome.” Science (New York, N.Y.) 350 (6264): 1096. https://doi.org/10.1126/science.aac7041.

Wang, Yi, Yanling Niu, and Bing Li. 2015. “Balancing Acts of SRI and an Auto-Inhibitory Domain Specify Set2 Function at Transcribed Chromatin.” Nucleic Acids Research 43 (10): 4881–92. https://doi.org/10.1093/nar/gkv393.

Wessels, Hans-Hermann, Alejandro Méndez-Mancilla, Xinyi Guo, Mateusz Legut, Zharko Daniloski, and Neville E. Sanjana. 2020. “Massively Parallel Cas13 Screens Reveal Principles for Guide RNA Design.” Nature Biotechnology 38 (6): 722–27. https://doi.org/10.1038/s41587-020-0456-9.

Yoh, Sunnie M., Joseph S. Lucas, and Katherine A. Jones. 2008. “The Iws1:Spt6:CTD Complex Controls Cotranscriptional MRNA Biosynthesis and HYPB/Setd2-Mediated Histone H3K36 Methylation.” Genes & Development 22 (24): 3422–34. https://doi.org/10.1101/gad.1720008.

Youdell, Michael L., Kelby O. Kizer, Elena Kisseleva-Romanova, Stephen M. Fuchs, Eris Duro, Brian D. Strahl, and Jane Mellor. 2008. “Roles for Ctk1 and Spt6 in Regulating the Different Methylation States of Histone H3 Lysine 36.” Molecular and Cellular Biology 28 (16): 4915–26. https://doi.org/10.1128/MCB.00001-08.

Yuan, Huairui, Ying Han, Xuege Wang, Ni Li, Qiuli Liu, Yuye Yin, Hanling Wang, et al. 2020. “SETD2 Restricts Prostate Cancer Metastasis by Integrating EZH2 and AMPK Signaling Pathways.” Cancer Cell 38 (3): 350-365.e7. https://doi.org/10.1016/j.ccell.2020.05.022.

Yuan, Wen, Jingwei Xie, Chengzu Long, Hediye Erdjument-Bromage, Xiaojun Ding, Yong Zheng, Paul Tempst, She Chen, Bing Zhu, and Danny Reinberg. 2009. “Heterogeneous Nuclear Ribonucleoprotein L Is a Subunit of Human KMT3a/Set2 Complex Required for H3 Lys-36 Trimethylation Activity in Vivo.” The Journal of Biological Chemistry 284 (23): 15701–7. https://doi.org/10.1074/jbc.M808431200.

Zaghi, Mattia, Vania Broccoli, and Alessandro Sessa. 2020. “H3K36 Methylation in Neural Development and Associated Diseases.” Frontiers in Genetics 10: 1291. https://doi.org/10.3389/fgene.2019.01291.

Zhang, Jinghui, Li Ding, Linda Holmfeldt, Gang Wu, Sue L. Heatley, Debbie Payne-Turner, John Easton, et al. 2012. “The Genetic Basis of Early T-Cell Precursor Acute Lymphoblastic Leukaemia.” Nature 481 (7380): 157–63. https://doi.org/10.1038/nature10725.

Zhang, Yinglu, Chun-Min Shan, Jiyong Wang, Kehan Bao, Liang Tong, and Songtao Jia. 2017. “Molecular Basis for the Role of Oncogenic Histone Mutations in Modulating H3K36 Methylation.” Scientific Reports 7 (1): 43906. https://doi.org/10.1038/srep43906.

Zhu, Xiaofan, Fuhong He, Huimin Zeng, Shaoping Ling, Aili Chen, Yaqin Wang, Xiaomei Yan, et al. 2014. “Identification of Functional Cooperative Mutations of SETD2 in Human Acute Leukemia.” Nature Genetics 46 (3): 287–93. https://doi.org/10.1038/ng.2894.

